# Multi-donor human cortical Chimeroids reveal individual susceptibility to neurotoxic triggers

**DOI:** 10.1101/2023.10.05.558331

**Authors:** Noelia Antón Bolaños, Irene Faravelli, Tyler Faits, Sophia Andreadis, Sebastiano Trattaro, Rahel Kastli, Xian Adiconis, Daniela J. Di Bella, Matthew Tegtmeyer, Ralda Nehme, Joshua Z. Levin, Aviv Regev, Paola Arlotta

## Abstract

Inter-individual genetic variation affects the susceptibility to and progression of many diseases. Efforts to study the molecular mechanisms mediating the impact of human genetic variation on normal development and disease phenotypes are limited, however, by the paucity of faithful cellular human models, and the difficulty of scaling current systems to represent multiple individuals. Here, we present human brain “Chimeroids”, a highly reproducible, multi-donor human brain cortical organoid model generated by the co-development of cells from a panel of individual donors in a single organoid, while maintaining fidelity to endogenous tissue. By reaggregating cells from multiple single-donor organoids at the neural stem or committed progenitor cell stage, we generate Chimeroids in which each donor produces all cell lineages of the cerebral cortex, even when using pluripotent stem cell lines with notable growth biases. We leveraged Chimeroids to investigate inter-individual variation in susceptibility to neurotoxic stressors that exhibit high clinical phenotypic variability: ethanol and the anti-epileptic drug valproic acid. Individual donors varied in both the penetrance of the effect on target cell types, and the molecular phenotype within each affected cell type. Our results show that human genetic background may be an important mediator of neurotoxin susceptibility and introduce Chimeroids as a scalable system for high-throughput investigation of the contribution of human genetic variation to brain development and disease.

## Introduction

Genetic variation plays a crucial role in differential susceptibility to disease triggers (Paulsen et al., 2022; Pizzo et al., 2019), such that different individuals show heterogeneous responses to disease risk factors. However, while the effects of human genetic variation can be detected in *in vitro* systems (Ford et al., 2022; Germain & Testa, 2017; Paulsen et al., 2022; Tegtmeyer et al., 2023), exploring the mechanisms that mediate the effect of common and rare genetic variants on physiological phenotypes and that drive differential responses between individuals is challenging. Investigation of these mechanisms is hampered by the limited availability of human experimental models that are both faithful to human endogenous biology and can be scaled across many individuals.

In the brain, pioneering studies have adapted pluripotent stem cell (PSC)-derived 2-dimensional (2D) neuronal cultures to produce population-scale models comprising multiple donor cell lines of different genetic backgrounds (Cederquist et al., 2020; Cuomo et al., 2020; Jerber et al., 2021; Limone et al., 2023; Mitchell et al., 2020; Wells et al., 2023). However, 2D cultures typically produce a limited spectrum of cell types that does not reflect the diversity of the endogenous brain. Three-dimensional (3D) cell culture models, such as human brain organoids, can more closely model the cellular complexity and developmental events of the endogenous brain, and thus may be used to test the effects of inter-individual variation across a wider range of cell types and developmental processes. However, efforts to extend the 2D “village-in-a-dish” approach (Mitchell et al., 2020; Wells et al., 2023) to 3D organoid cultures have been limited, because differences in growth rate and differentiation biases between PSC lines lead to unbalanced donor representation in multiplexed PSC differentiation systems, such that certain lines take over the cultures or are eliminated. Prior selection of PSC lines with matched growth rates can ameliorate the problem (Wells et al., 2023), but limits experimental design, in that many PSC lines cannot be used. Such biases are particularly problematic for organoid systems, where multiple cell types co-develop over long time periods, exacerbating the effect of growth or differentiation biases (Jerber et al., 2021).

Here, we addressed this challenge by developing 3D multi-donor Chimeroids: a highly-reproducible, human cortical organoid model that preserves balanced representation of cell types derived from panels of donors, without the need for PSC pre-selection. By fine-tuning the developmental stage when donor line multiplexing occurs, we diminished differences in growth biases among donors in the resulting chimeroids. We used this model to measure inter-individual susceptibility to developmental exposure to two neurotoxic stressors: ethanol (EtOH, associated with Fetal Alcoholic Syndrome (Wozniak et al., 2018)) and the anti-epileptic drug valproic acid (VPA, associated with increased risk of Autism Spectrum Disorders (Bjørk et al., 2022; Christensen et al., 2013)). EtOH and VPA exposure resulted in changes across multiple cell types within Chimeroids, and, importantly, Chimeroids captured donor-specific differences in response. Overall, our findings show that inter-individual variation is a significant source of susceptibility to neurotoxic triggers, and that Chimeroids can be used as a scalable platform to measure variation in biological response in brain tissue of different human individuals.

### Brain Chimeroids produce cortical cell diversity balanced across individual donors

To develop a human brain system that can model variation in responses by different individuals to disease risk factors, such as genetic variants or neurotoxic insults, we devised a strategy for generating multi-donor human brain organoids, in which each cell type would be produced from multiple donor lines within a single brain organoid.

Because human PSCs from multiple donors can be grown and differentiated together in 2D cultures (Cederquist et al., 2020; Cuomo et al., 2020; Jerber et al., 2021; Limone et al., 2023; Mitchell et al., 2020; Neavin et al., 2023; Villa et al., 2022; Warren & Cowan, 2018; Wells et al., 2023), we first attempted to generate Chimeroids by mixing 4 to 5 lines in equal ratios at the PSC stage (referred to here as PSC-Chimeroids; **Fig. 1A-C** and **Suppl. Fig. 1A-F**, **Suppl. Table 1**). We used the following lines: H1 (male, human embryonic stem cell [hESC]), Mito210 (male, induced pluripotent stem cell [iPSC], control with no familial history of neurological disorders), PGP1 (male, iPSC, control), CW50037 (hereafter ‘CW’; female, iPSC, control with no familial history of neurological disorders), GM08330 (hereafter ‘GM’; male, iPSCs, clinically-unaffected with familial history of relevant psychiatric illness), and 11a (male, iPSC, control).

**Figure 1.**
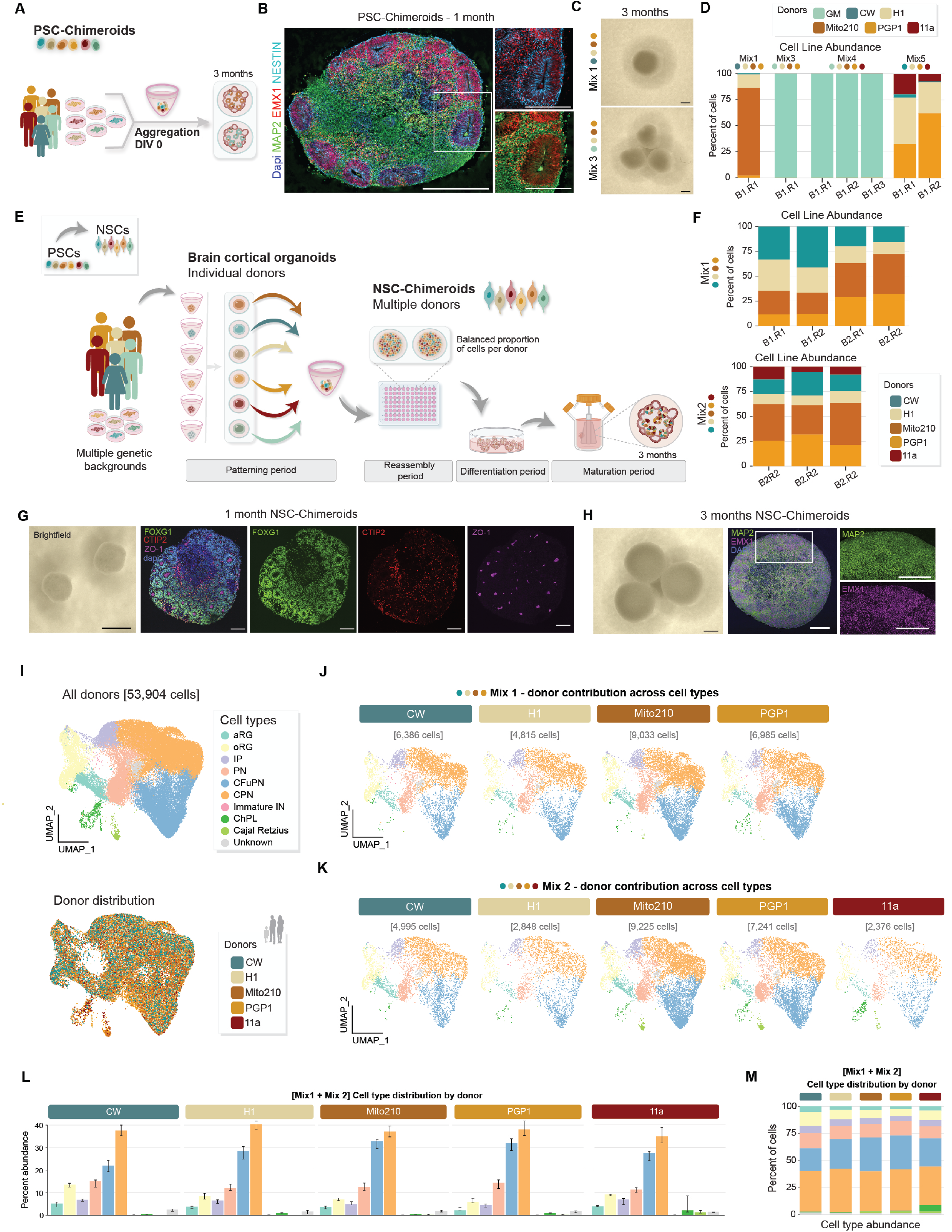
NSC-Chimeroids maintain donor representation across cell types. **A,** Schematic of the PSC-Chimeroids protocol. PSCs: pluripotent stem cells, DIV: days *in vitro*. **B**, Immunolabelling of MAP2, EMX1 and NESTIN of cortical PSC-Chimeroids at 1 month (mo), whole organoid and zoom-in images. Scale bar: 500 μm and 250 μm respectively. **c**, Brightfield images of cortical PSC-Chimeroids at 3 mo. Scale bar: 1 mm. **D**, 3-mo PSC-Chimeroids analyzed by single-cell RNA-seq and demultiplexed using demuxlet show unequal distribution of donor lines; donor is indicated by colors. *B*= batch and *R*= biological replicate. **e,** Schematic of the NSC-Chimeroids protocol. NSCs: Neural Stem Cells. **F**, Donor NSC-Chimeroids demultiplexed after single-cell RNA-seq at 3 mo. **G**, Left panel: brightfield images of cortical NSC-Chimeroids at 1 mo. Scale bar: 1 mm. Right panel: immunolabelling of 1-mo cortical NSC-Chimeroids showing cortical progenitors (FOXG1), corticofugal pyramidal neurons (CTIP2), and tight junctions formed by neural progenitors (ZO-1). Scale bar: 100 μm. **H**, Brightfield images of 3-mo cortical NSC-Chimeroids (n=3; left panel). Scale bar: 1 mm. Immunolabelling 3-mo cortical NSC-Chimeroids showing cortical progenitors (EMX1) and the neuronal dendritic marker MAP2. Scale bar: 500 μm. **i**, UMAP of integrated NSC-Chimeroids, color-coded by annotated cell type (upper panel) and donor line, and quantification of donor contribution with demuxlet (lower panel). **J-K**. UMAPs split by donor for two different mixes (Mix 1, 4 donors and Mix 2, 5 donors). **L**, Bar plots of cell-type proportions in NSC-Chimeroids, demultiplexed by donor. Whiskers show upper and lower quartile values across replicates (n=7 for CW, H1, Mito210, and PGP1; n=3 for 11a). **M**, Stacked bar plots of cell-type proportion for each donor within Mix 1 and Mix 2.

Mixes of PSC lines were aggregated in 96-well plates to form embryoid bodies (EBs); then, following our previously-published protocol (Velasco et al., 2019), EBs were patterned for 15-18 days to induce dorsal forebrain fates, and nascent organoids were then switched to CDMII media and placed in dynamic (agitated) culture to allow maturation (**Methods**). We quantitatively determined the contribution of individual donor lines to each organoid at different stages of development using Census-seq (Mitchell et al., 2020), which uses low-coverage whole genome sequencing (WGS) to determine donor proportion.

At 1 month *in vitro* (1 mo), PSC-Chimeroids showed unbalanced contribution of the individual donors, which became more pronounced upon further culture, at 3 mo (**Fig.1D** and **Suppl. Fig. 1B**). We tested four different combinations of donors, from distinct experiments, at different time points, and observed unbalanced growth in all cases (**Fig. 1D** and **Suppl. Fig. 1B**). Specifically, we analyzed: Mix 1 (4 lines: CW, H1, Mito210, PGP1), Mix 3 (4 lines: GM, H1, Mito210, PGP1), Mix 4 (5 lines: GM, H1, Mito210, PGP1, 11a) and Mix 5 (4 lines: CW, H1, PGP1, 11a). PSC-Chimeroids that included GM (Mix 3 and Mix 4) had pronounced overgrowth of this line, which almost completely took over the Chimeroids by 2 mo (**Suppl. Fig. 1B**). At 23 days *in vitro* (DIV23), 54% of cells in Mix4 PSC-Chimeroids were GM-derived (**Suppl. Fig. 1B**)=, and single-cell RNA-seq (scRNA-seq) on Chimeroids from this mix followed by supervised donor assignment by genetic variants (Kang et al., 2018) (**Methods**) showed that at 3 mo the GM line contributed almost all (>99%) of the cells in the PSC-Chimeroids (**Fig. 1D**). In the other mixes (without GM), the Mito210 cell line was disproportionately overrepresented (84% of the cells in Mix 1 PSC-Chimeroids at 3 mo, **Fig. 1D**). In Mix 5 PSC-Chimeroids, which had neither GM nor Mito210, one line (CW) was almost eliminated (<3% of the Chimeroid cells at 3 mo, **Fig. 1D** and **Suppl. Fig. 1D**). We conclude that mixing different PSC lines at the initiation of EB formation leads to vastly disproportionate representation of individual donors, a finding that is consistent with previous observations in a dopaminergic neuron model(Jerber et al., 2021). Despite these differences between lines, when multiple donors maintained significant representation (Mix 5), all retained donors were able to contribute to all identifiable cell types in a largely balanced manner, indicating that the Chimeroid approach does not *per se* alter cell fate potential in individual donors (**Fig.1B** and **Suppl. Fig.1D-F**).

PSCs are highly proliferative and plastic in fate potential, and small differences in these properties early on during organoid development may lead to disproportionate representation at later times. To constrain such inter-line biases, we hypothesized that mixing different donor lines after PSCs have undergone neural fate commitment, when they have reduced cell growth and more restricted fate potential, would reduce variability in terminal donor contribution. We therefore modified our Chimeroid protocol such that donor mixing would occur after completion of neural patterning, at the neural stem cell (NSC) stage (referred to hereafter as NSC-Chimeroids) (**Fig. 1E**).

To create NSC-Chimeroids, we first generated single-donor EBs, and patterned them with small molecules until DIV15-18(Velasco et al., 2019). We dissociated nascent single-donor organoids from 4 to 5 individual lines to single cells, and mixed donor cells at equal proportions (DIV 0, **Suppl. Fig. 2A**). We then seeded the pooled cells in 96-well plates and allowed them to re-aggregate for 2 days before transfer to dynamic culture (**Fig. 1E** and **Methods**). At 20 days after aggregation, NSC-Chimeroids had developed ‘rosettes’, prospective ventricular zone-like (VZ-like) structures composed of cells expressing the telencephalic marker FOXG1, surrounded by nascent corticofugal projection neurons (CTIP2^+^); the rosette centers were lined with ZO-1, a tight junction protein present in the endfeet of apical radial glia (aRG), indicating correctly-polarized epithelium (**Fig. 1G**).

Quantification by Census-seq (n=36 individual Chimeroids) and scRNA-seq (n=7 individual Chimeroids, 53,904 cells) showed that NSC-Chimeroids maintained substantially more balanced contribution of each donor line, in each of 4 different mixes (**Fig. 1F** and **Suppl. Fig. 2A**). Notably, Mix 1, 3 and 4, which in PSC-Chimeroids were dominated (85% to 99%) by a single donor, retained contribution of all donors in NSC-Chimeroids. Thus, mixing donor lines at the NSC stage improves retention of individual donors, compared to mixing at the PSC stage.

At 3 mo, NSC-Chimeroids showed proper expression of dorsal forebrain markers, at the protein level (**Fig. 1H**). ScRNA-seq of Mix 1 and Mix 2 NSC-Chimeroids at 3 mo (n=7 individual Chimeroids, 53,904 cells) showed that both mixes generated the full compendium of cell types expected from single-donor dorsal forebrain organoids produced by our original protocol (Velasco et al., 2019) (**Fig. 1I-K**). Importantly, each donor contributed to the generation of each cell type (**Fig. 1L-M**, **Suppl. Fig. 2B-E**, **Suppl. Fig. 3**).

### Chimeroids can control donor biases in proliferation

To investigate the potential of Chimeroids to buffer against donor variation in growth rates, we challenged the system by including one iPSC line with a dramatic proliferative advantage (GM) (**Fig. 1D**). We created multi-donor NSC-Chimeroids from two additional mixes of 4 or 5 lines: Mix 3 (GM, H1, Mito210, and PGP1) and Mix 4 (GM, H1, Mito210, PGP1, and 11a) (**Suppl. Table 1**). Although incorporating GM skewed the proportions of donors in NSC-Chimeroids (GM contributing 76-95% of cells; **Fig. 2A-C**), at 3 months all donors were represented and able to generate all the cell types expected at this stage (Uzquiano et al., 2022) (**Fig. 2C-E**). GM produced a higher proportion of neural precursor cells (mean=31.0%, sd=3.28%) compared to other donors (mean=13.8%, sd=6.4%), showing a particular expansion of oRGs (GM: mean=16.1%, sd=0.6%; other donors: mean=6.8%, sd=2.6%; **Fig. 2C**), thus pointing at possible reasons for the growth advantage of the GM line. Consistent with this, pseudotime analysis suggested an increased proportion of cells predicted to occupy earlier developmental stages in GM-derived cells compared to cells from other donors (**Fig. 2F**).

**Figure 2.**
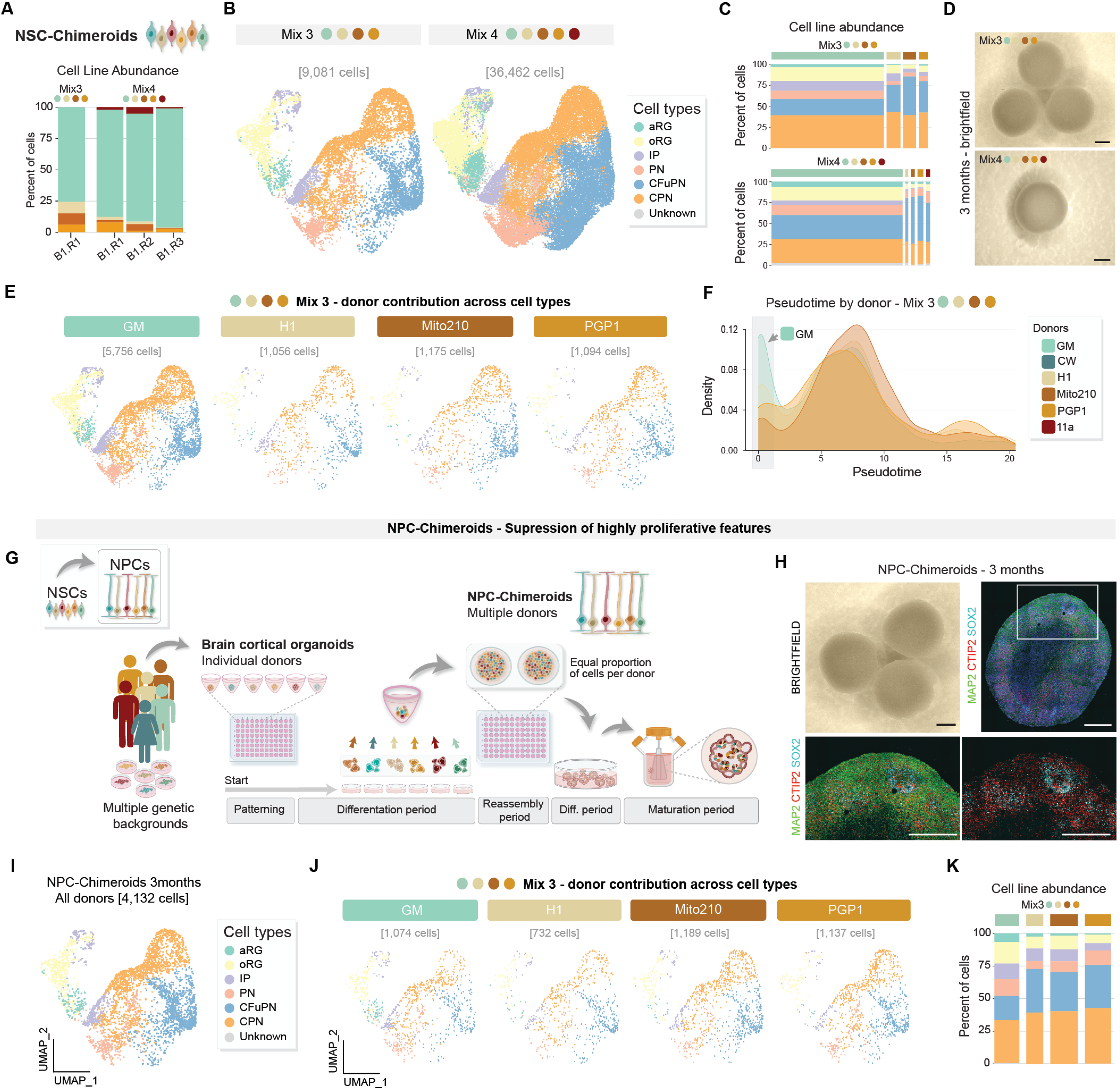
NSC- and NPC-Chimeroids control hyperproliferative states of pluripotent stem cells. **A,** 3-mo NSC-Chimeroids analyzed by single-cell RNA-seq and demultiplexed by donor: Mix 3 and Mix 4 include the highly proliferative donor GM. **B,** 3-mo NSC-Chimeroid UMAPs color-coded by annotated cell type for Mix 3 (4 donors) and Mix 4 (5 donors). **C,** Bar plot with cell type and donor composition; the width of each bar corresponds to the donor proportion. **D,** Brightfield images of 3-mo cortical NSC-Chimeroids. Scale bar: 1 mm. **E,** UMAPs split by donor for Mix 3 (4 donors). **F,** Density plot of pseudotime by cell line, calculated using Monocle3 with all Cycling cells as roots. **G**. Schematic of protocol for NPC-Chimeroids. Differentiation= diff. **H**, Upper-left panel: brightfield of 3-mo NSC-Chimeroid. Scale bar: 1 mm. Immunohistochemistry of 3-mo NPC-Chimeroids showing SOX2 (progenitors), CTIP2 (cortical neurons) and MAP2 (dendrites). Scale bar: 500 μm. **I-J.** UMAP representation of NPC-Chimeroids integrated (**I**) and split by donor (**J**). **K**, Barplots of cell type and donor composition demonstrate balanced growth and proper cortical development; the width of each bar corresponds to the donor proportion.

Following this observation, we hypothesized that mixing progenitors at even later, less-proliferative stages of differentiation and fate commitment might further improve donor growth biases. We therefore generated Chimeroids from neural progenitor cells (NPCs), by dissociating and mixing cells at a later stage of the organoid protocol, at DIV23-25 (Mix 3: GM, Mito210, H1, PGP1), and profiled these NPC-Chimeroids by scRNA-seq (1 batch, n=2, 4,132 cells) at DIV90 (**Fig. 2G-K**). NPC-Chimeroids achieved the expected cell type composition (**Fig. 2I**) and, importantly, had more balanced donor contribution, even in the presence of the GM line that overwhelmingly dominated the PSC-Chimeroids. Specifically, at DIV90, GM contributed 17-31% of cells in NPC-Chimeroids (**Fig. 2K**) vs. 76-95% and >99% of cells in NSC- and PSC-Chimeroids, respectively. Notably, GM still showed a higher proportion of proliferative cell types (aRG, oRG, and IP) compared to cells from other donors (34.4%-35.5% in GM vs 11.0%-23.8%; binomial mixed effects model, p<10^-15^. **Fig. 2K**), indicating that the Chimeroid protocol preserves intrinsic differences between donors. Overall, Chimeroid technology allows using neural progenitors at different stages of development as starting cell types for aggregation, and performing donor mixing at later stages of organoid development can overcome substantial donor-specific growth advantages, without requiring prior knowledge of line-specific properties.

### Cellular diversification in NSC- and NPC-Chimeroids is comparable to single-donor organoids

To understand if the cell types produced in Chimeroids are comparable to those produced in single-donor organoids, we compared the cell compositions produced by the same donor lines in the different models. To control for any effect of dissociation and, we generated single-donor NSC-Chimeroids by dissociation and reaggregation of nascent organoids from a single donor line. We profiled 53,904 cells from individual NSC-Chimeroids (from two different mixes, Mix 1 and Mix 2), 4,132 cells from NPC-Chimeroids, and 35,180 cells from single-donor Chimeroids, all at 3 mo, and compared them to our published single-cell atlas of cortical organoid development (Uzquiano et al., 2022).

All multidonor Chimeroids contained the same cell type subsets present in 3 mo single-donor organoids from our published reference atlas, including apical radial glia (aRG), outer radial glia (oRG), intermediate precursor cells (IPC), and corticofugal (CFuPN) and callosal (CPN) projection neurons (**Fig. 3A-C**). Moreover, the expression profiles of 19 canonical cell-type marker genes for key cell populations were highly concordant between Chimeroids and single-donor organoids from the atlas (Pearson correlation coefficients for normalized marker gene expression between corresponding cell types ranging 0.93-0.98; **Fig. 3B and Suppl. Table 2**), and between corresponding cell types of Chimeroids and fetal cortex (Pearson correlation coefficients ranging 0.81-0.91, **Fig. 3C**). The correlation between expression of these marker genes in NSC-Chimeroids and fetal cortex was not significantly different from the correlations between single-donor atlas organoids and fetal cortex (paired Wilcoxson test, p=0.19) or between single-donor NSC-Chimeroids and fetal cortex (p=0.06), indicating that Chimeroid cell types demonstrate equally high similarity to endogenous cell types. There was also a high degree of global similarity between expression profiles in corresponding cell types in the NSC-Chimeroids compared to both our organoid atlas (Uzquiano et al., 2022) and endogenous fetal tissue (rank-rank hypergeometric overlap (RR-HO) (Cahill et al., 2018), **Methods**, **Fig. 3D**, **Suppl. Fig. 4-5**). Finally, reference label transfer (Stuart et al., 2019) from NSC-Chimeroids to cell types from two reference datasets, endogenous human fetal cortex and single-donor organoids, or vice versa (**Methods**), showed that Chimeroid cells were predominantly assigned to the expected reference cell type in both datasets, indicating high similarity in expression profiles (**Fig. 3E**).

**Figure 3.**
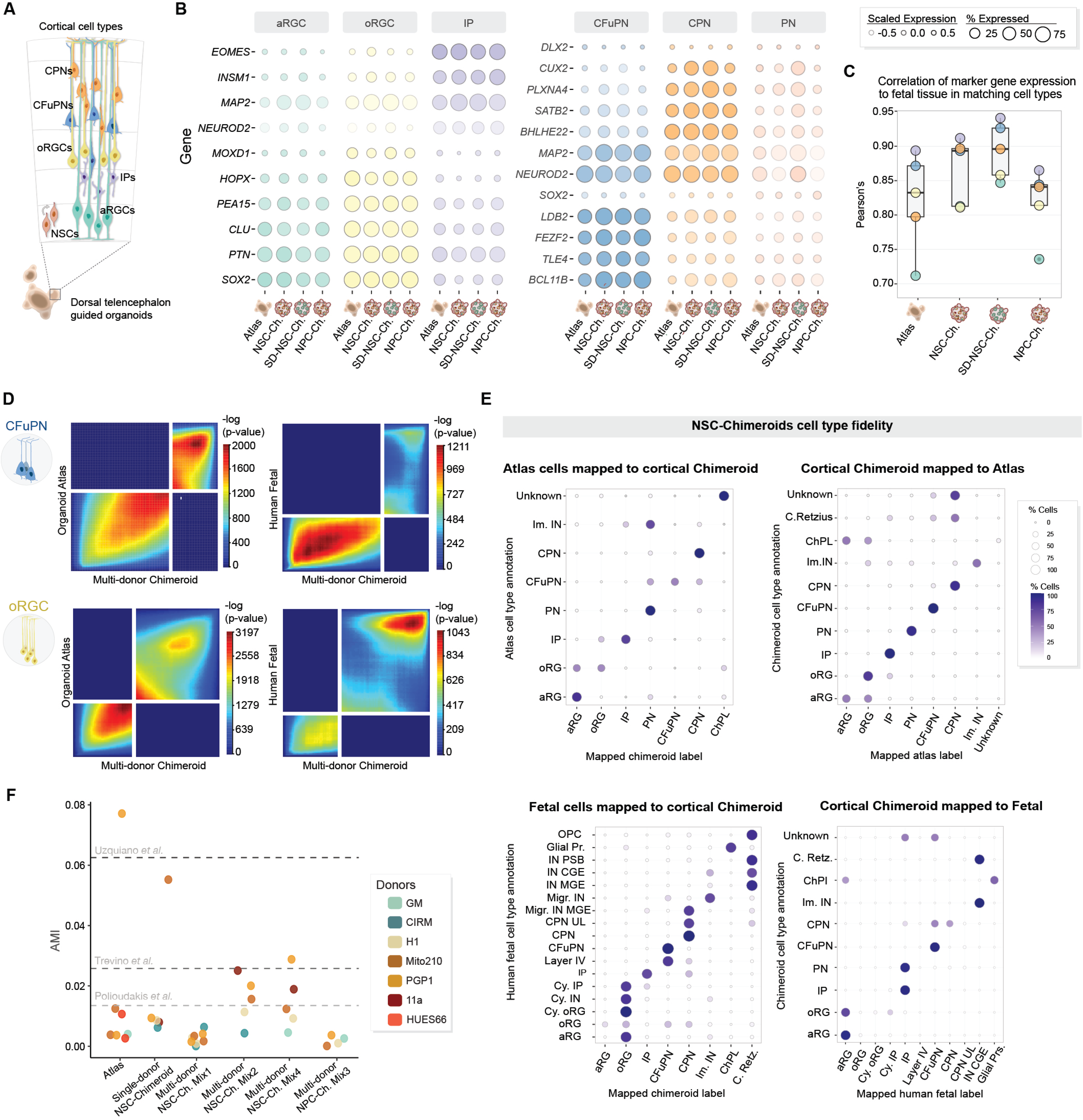
Chimeroids are reproducible and display proper cell-type composition. **A,** Schematic of cell types in the developing cortex. **B**, Dot plots showing expression of marker genes across common cell types in a previously published atlas of cortical organoid development (‘atlas’) (Uzquiano et al., 2022), NSC-Chimeroids, single-donor (SD) NSC-Chimeroids, and NPC-Chimeroids. Point size indicates the percentage of cells which express each marker gene, and shade represents average expression values. Color indicates annotated cell type. **C**, Pearson correlation coefficients for normalized marker gene expression between human fetal cortical cell types and corresponding cell types in atlas organoids, NSC-Chimeroids, single-donor NSC-Chimeroids, and NPC-Chimeroids. **D**, Rank-Rank Hypergeometric Overlap (RR-HO) plots comparing the expression signatures of CFuPN and oRGC cell types (upper and lower panels respectively) in multi-donor Chimeroids to their counterparts in atlas cortical organoids or endogenous fetal tissue. The horizontal axes represent genes significantly differentially expressed in a given Chimeroid cell type compared to all other Chimeroid cells, ranked from most upregulated to most downregulated; the vertical axes represent similarly ranked lists of differentially-expressed genes for atlas organoid or fetal cells. Color at a given position represents the significance (negative log p-value) of the overlap of the gene lists up to that point, as calculated by Fisher’s exact tests. High significance (i.e., red color) in the lower left and upper right quadrants indicates strong concordance between the expression profiles which define the corresponding cell types. **E,** Left, cell type annotations assigned to Chimeroid cells using two different reference datasets: an atlas of human cortical organoid cells (top left), and endogenous human fetal cortex cells (lower left). Right, the converse: cell type annotations using Chimeroid cells as a reference, mapped onto cells from the cortical organoid atlas (top right) and from fetal tissue (lower right). The vertical axis groups cells by manually-annotated cell type (based on marker gene expression and clustering in the query dataset). The horizontal axis groups cells by the transferred label from the reference dataset. Point size and color indicates the percentage of cells which were assigned a given label. **F**, Adjusted Mutual Information (AMI) values within batches between cell types and individual cortical organoids from the atlas, individual single-donor NSC-Chimeroids, and individual donors within each Mix 1, 2, 4 NSC-Chimeroid and Mix 3 NPC-Chimeroids. Lower values indicate lower variability in cell type composition between replicates. Dotted lines represent AMI values between replicates within three datasets of endogenous human fetal cortex (Polioudakis et al., 2019; Trevino et al., 2021; Uzquiano et al., 2022).

To assess Chimeroid-to-Chimeroid variability in cell-type composition, we calculated the adjusted mutual information (AMI) (*Information Theoretic Measures for Clusterings Comparison: Variants, Properties, Normalization and Correction for Chance*, n.d.) of cell-type abundances between replicates within each batch of organoids from the atlas, as well as between replicates for each donor in each batch of single-donor and multi-donor NSC-Chimeroids. AMI scores for multi-donor NSC-Chimeroids or for single-donor Chimeroids were not significantly different from those for atlas organoids (Wilcoxon rank-sum test, p=0.98 and p=0.53, respectively) (**Fig. 3F**), indicating that Chimeroids retain high chimeroid-to-chimeroid reproducibility comparable to that of our established cortical organoid protocol (**Fig. 3F**). Notably, 3-mo Chimeroid replicates had AMI scores comparable to those of individual human fetal brains profiled in published datasets (Polioudakis et al., 2019; Trevino et al., 2021; Uzquiano et al., 2022) (**Fig. 3F**, dashed lines). Interestingly, within individual multi-donor Chimeroids, the variability in cell-type composition between donors was low (mean AMI=0.0096), and not different from the organoid-to-organoid AMI found in single-donor cortical organoids (Wilcoxon test p=0.9), indicating that donor-specific differences in cell type composition are consistent across individual Chimeroids.

Overall, these data show that the Chimeroid model is reproducible between replicates of the same mixture of donors and is concordant with our original single-donor organoid model. Multi-donor Chimeroids also retain the same degree of cellular complexity and molecular resemblance to endogenous human fetal tissue as the original cortical organoid model (Uzquiano et al., 2022; Velasco et al., 2019).

### Ethanol and valproic acid induce cell type-specific effects in Chimeroids

To demonstrate the utility of the Chimeroid model to investigate inter-individual variability in response to perturbation, we tested the effects of two neurotoxic triggers that cause neurodevelopmental abnormalities: ethanol (EtOH) and valproic acid (VPA). Fetal exposure to EtOH in humans can result in Fetal Alcohol Spectrum Disorders (FASD) (Alfonso-Loeches & Guerri, 2011; Arzua et al., 2020; Carpita et al., 2022; Charness, 2022; Eberhart & Parnell, 2016; Granato & Dering, 2018; Marguet et al., 2022; Streissguth & Dehaene, 1993; Sulik et al., 1981; Wozniak et al., 2018), with high inter-individual variation in clinical phenotype(Eberhart & Parnell, 2016). Valproic acid (VPA), used to treat epilepsy and psychiatric disorders, is associated with augmented risk of developing autism related disorders (ASD) in offspring when administered during pregnancy (Bjørk et al., 2022; Christensen et al., 2013).

We generated NSC-Chimeroids from two different mixes, Mix 1 and Mix 2 (**Suppl. Table 1**), and treated them with either EtOH or VPA for 30 days. To model physiological fluctuations typical of alcohol exposure during pregnancy, we administered 50 μM of EtOH to the culture media every two days, from DIV45 to DIV75, with plates sealed for 24 hours after EtOH addition to avoid evaporation (Arzua et al., 2020) (**Fig. 4A**, top). To mimic environmental exposure to VPA, we added VPA at clinical concentrations (0.7 mM) directly to the culture media two times per week, from DIV45 to DIV75 (**Fig. 4A**, bottom). Both treatments were stopped at DIV75, and Chimeroids were cultured until DIV90. Finally, we profiled the treated 3-mo NSC-Chimeroids from two donor mixes by scRNA-seq. We jointly annotated the cells from both treatment groups, as well as the untreated Mix 1 and Mix 2 NSC-Chimeroids controls (above), by combining all VPA, EtOH, and control data, followed by clustering (**Methods**), and annotated cell clusters by expression of canonical marker genes and examination of genes upregulated in each cluster (Mix1 94,009 cells from n=12 Chimeroids, Mix2 88,406 cells from n=9 Chimeroids; **Fig. 4B-E**, **Suppl. Fig. 6**).

**Figure 4.**
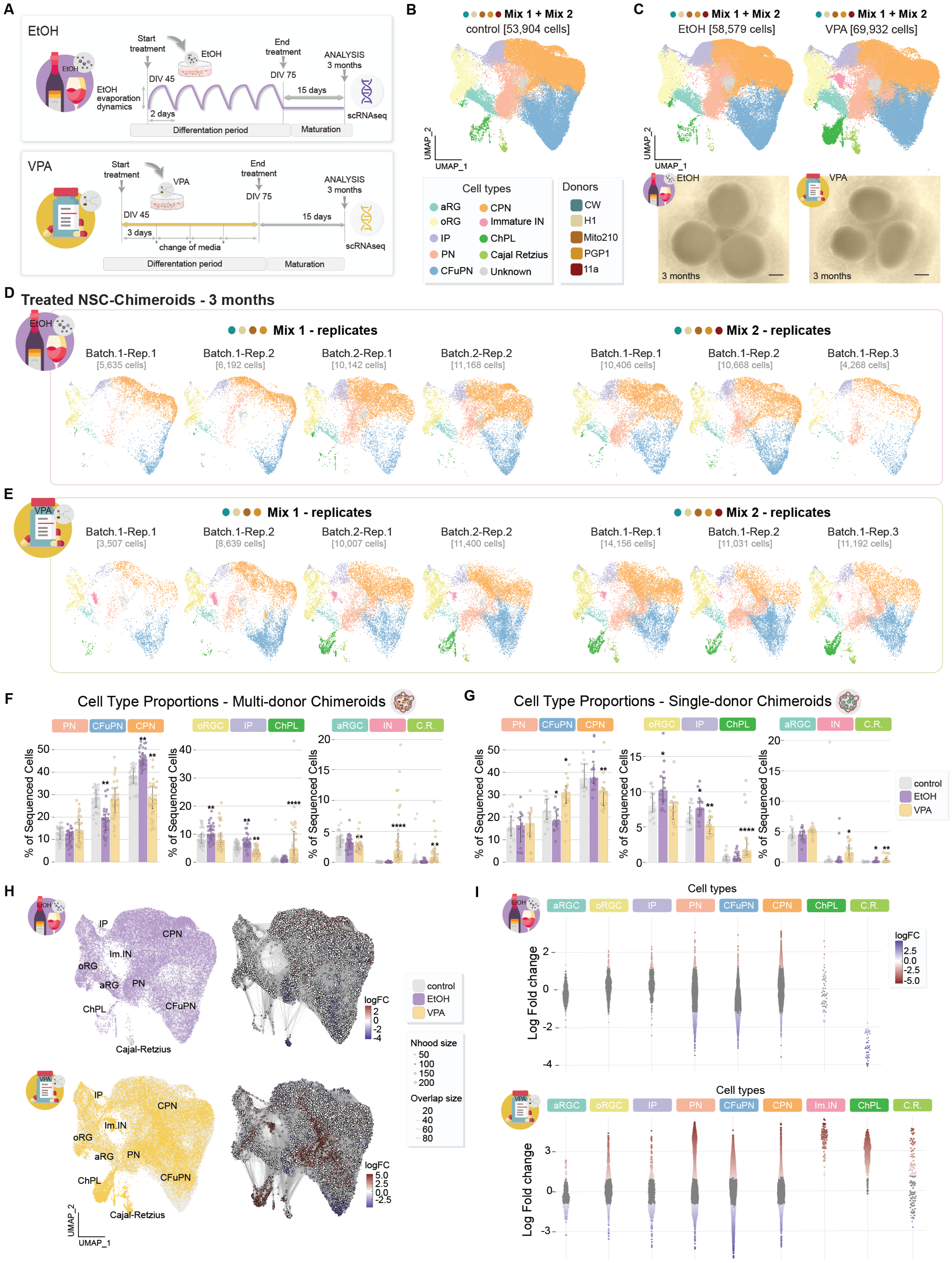
NSC-Chimeroids model treatment-specific alterations. **A.** Schematic of the EtOH and VPA experimental design. **B-C**, Integrated UMAPs colored by annotated cell type for control (**B**) and treated (**C**) conditions, with corresponding brightfield images of 3-mo NSC-Chimeroids. Scale bar: 1 mm. **D-E**, UMAPs of EtOH (**D**) and VPA (**E**) treated NSC-Chimeroids, split by mixes and replicates. **F**, Bar plots of cell-type proportions in control multi-donor NSC-Chimeroids in control, (grey) and treated with EtOH (purple) or VPA (yellow). Whiskers show upper and lower quartiles. Plot is split into three facets to allow rescaling of the y-axis (indicating percent abundance). Significance of differences between treated and control Chimeroids was calculated using a negative binomial generalized linear model. FDR-adjusted p-values: * - <0.1; ** - <0.01; *** - <0.001. **G**. Changes in cell-type proportions in single-donor Chimeroids from the corresponding lines, as in f. **H**, Left: UMAP showing the localization of cells from control multi-donor Chimeroids (grey) and cells from multi-donor Chimeroids treated with VPA (yellow) and EtOH (violet). Right: UMAP showing overlapping neighborhoods of cells, as calculated using Milo. Red and blue colors indicate neighborhoods with significant enrichment for EtOH and VPA-treated cells or control cells, respectively. Point size indicates the number of cells in a neighborhood, and edge thickness indicates the number of cells shared between pairs of neighborhoods. **I**, Beeswarm plot showing shifts in the composition of neighborhoods of cells in response to EtOH and VPA treatment, grouped by the cell-type identity of those neighborhoods. Each point represents a neighborhood of 50-200 cells with similar gene expression profiles. The vertical axis indicates the enrichment of EtOH and VPA-treated cells within a neighborhood, with positive log2 fold change values indicating more than expected EtOH or VPA-treated cells, and negative values indicating fewer than expected EtOH or VPA-treated cells. Neighborhoods are colored based on statistical significance of that enrichment: grey, not significantly different from random; red, significant over-enrichment; blue, significant under-enrichment. If most neighborhoods within a cell type collectively shift up or down, it implies an overall gain or loss, respectively, of that cell type in EtOH or VPA-treated Chimeroids. Cell types with neighborhoods that form long tails of both overenrichment and underenrichment are likely to have EtOH or VPA-induced changes in expression profile, without necessarily changing in abundance.

While control Chimeroids displayed the cellular composition previously observed in the single-donor organoid model at this age (**Fig. 4B**) (Uzquiano et al., 2022; Velasco et al., 2019), each treatment group showed distinct and significant alterations in cell-type composition (**Fig. 4F-G**). VPA treatment caused a dramatic increase in the proportion of immature GABAergic interneuron (FDR=9.4*10^-7^, negative binomial mixed-effects (NBME) model) and choroid plexus cells (FDR=3.0*10^-6^), even when accounting for batch, mix, and donor (**Methods**) (**Fig. 4f, Suppl. Fig. 7**), consistent with previous reports on VPA-treated organoid models (Meng et al., 2022). Interestingly, previous results from our group identified accelerated interneuron development as one of the convergent phenotypes of haploinsufficient loss-of-function mutations in different ASD risk genes (Paulsen et al., 2022), suggesting that this cell population may be particularly susceptible to neurodevelopmental risk factors. Other neuronal clusters also showed treatment-specific shifts in proportion: CPN proportions were increased by EtOH treatment (FDR=0.001, NBME model) and decreased in VPA-treated Chimeroids (FDR=0.01), and IP proportions were decreased by both treatments (FDR=0.0042 for VPA and 0.0038 for EtOH) (**Fig. 4F**).

Because statistical tests for change in proportions of individual cell types do not account for the interdependence between such changes (which must sum to zero), we confirmed these observations and assessed the effects of EtOH and VPA on global expression profiles using Milo (Dann et al., 2022), a method designed for testing differential cell abundance and expression variance in single-cell data without using cluster identity. Milo corroborated the increase in proportion of immature interneurons and choroid plexus cells, and decrease in proportion of aRGs, in VPA-treated Chimeroids, and the decrease in proportion of CFuPNs in EtOH-treated Chimeroids (**Fig. 4H-I**). Moreover, Milo highlighted global shifts in the expression landscape without overall change in discrete cell type proportions in oRGs, CFuPNs, and CPNs in VPA-treated Chimeroids, and in CPNs in EtOH-treated Chimeroids (**Fig. 4I**). We observed similar effects of VPA treatment on cell-type proportions in single-donor Chimeroids (**Fig. 4G**, **Suppl. Fig. 8**), although this analysis had less statistical power due to the smaller dataset size. Overall, VPA caused more expression changes than EtOH treatment, in line with its established role as an epigenetic effector (Göttlicher et al., 2001); in VPA-treated Chimeroids, 3,014 out of 8,848 neighborhoods were significantly altered, compared to only 520 out of 7,941 neighborhoods for EtOH-treated Chimeroids.

Overall, these findings demonstrate that VPA and EtOH cause treatment-specific alterations in cell composition and cell-intrinsic expression states that could be detected in multi-donor Chimeroids. Interestingly, VPA exposure, which is correlated with ASD (Bjørk et al., 2022; Christensen et al., 2013), induced an increase in the immature GABAergic interneuron population, reminiscent of a cellular phenotype induced by haploinsufficiency of several different ASD risk genes in single-donor cortical organoids (Paulsen et al., 2022). Moreover, similar phenotypic alterations could be detected in single-donor Chimeroids, thus corroborating the use of multi-donor Chimeroids to investigate treatment-specific responses.

### Chimeroids allow analysis of inter-individual variation in response to neurotoxins

We next asked whether we could detect donor-specific responses to EtOH and VPA treatment in Chimeroids. While each donor contributed to all cell types, regardless of perturbation (**Fig. 5A**), the overall proportion of Chimeroid cells derived from individual donors was differentially affected by VPA and EtOH, across two separate mixes (**Fig. 5B**). Specifically, the proportion of Chimeroid cells derived from the CW donor line was decreased by EtOH treatment (FDR=0.036, (NBME) model) and increased by VPA (FDR=6.5*10^-5^), indicating a treatment- and donor-specific effect of these two compounds (**Fig. 5B**).

**Figure 5.**
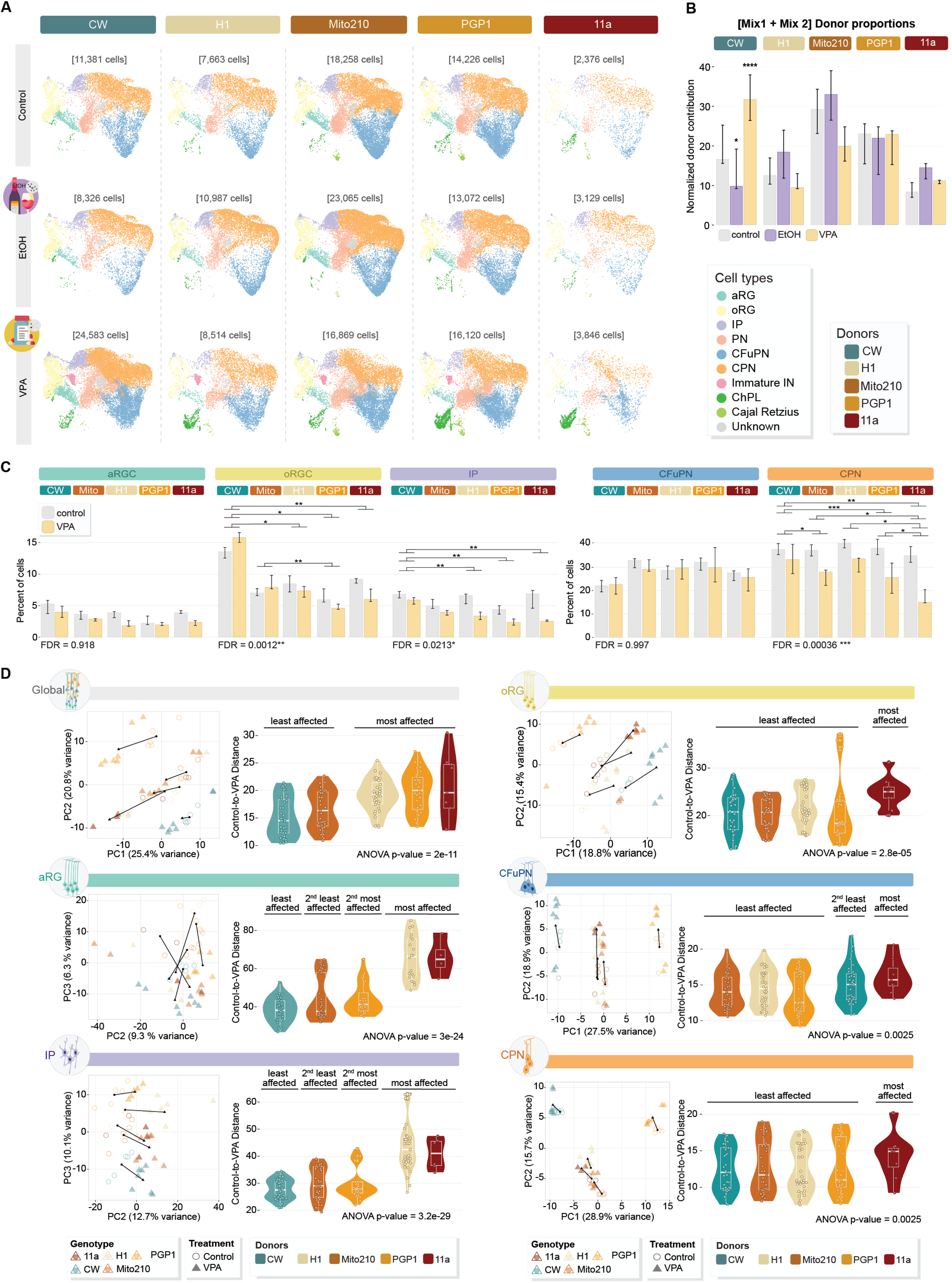
NSC-Chimeroids display differential donor-specific susceptibility. **A-C,** UMAPs of control, EtOH-, and VPA-treated Chimeroids split by donor and colored by cell type. **B**, Bar plot of median normalized donor proportions in control NSC-Chimeroids (grey) and treated with EtOH (purple) or VPA (yellow). The y-axis represents the ratio of the measured donor abundance in each Chimeroid to the expected abundance if all donors within a mix were equally represented (i.e., 25% in Mix 1 or 20% in Mix 2). Whiskers show upper and lower quartiles. Significance of differences between treated and control Chimeroids was calculated by applying a negative binomial generalized mixed effects model to the cell counts per donor. FDR-adjusted p-values: * - <0.1; ** - <0.01; *** - <0.001. **C**, Bar plot of VPA-induced changes in cell type proportions, split by donor. Whiskers show upper and lower quartiles. Significance was calculated using a negative binomial generalized mixed effects model, with treatment as a fixed effect, Chimeroid-of-origin as a random effect, donor as a random effect, and treatment-by-donor as a random interaction. FDR-adjusted p-values for the random interaction were calculated via likelihood ratio test against a model without the treatment-by-donor random interaction. For cell types with significant random interactions (oRG, IP, and CPN), post-hoc pairwise comparisons were performed between donors using the same models; significant pairwise differences in response to treatment are identified with brackets: * - <0.1; ** - < 0.01; *** - <0.001. **D**, Principal Component Analysis (PCA) performed on the gene expression matrix of different clusters from VPA-treated (triangles) and untreated (circles) NSC-Chimeroids, pseudobulked by donor (indicated by color) and replicate, both using all cells (top left) and subsetted by cell type. Data points separate in PC1 and PC2 by donor and treatment, except in the case of IP and aRG, where they separate in PC2 and PC3. Corresponding right panels: violin plots showing the pairwise Euclidean distance in PCA space between treated and untreated pseudobulked data points from each donor. Larger distances imply that VPA treatment induces larger changes in expression profiles. Significance calculated by Analysis of Variance (ANOVA), followed by Tukey post-hoc pairwise tests.

For a more granular analysis of donor-specific effect of each treatment, we tested whether, VPA treatment also differentially affected the proportion of individual cell types for each donor. We found that either the magnitude or the direction of change in representation induced by VPA was significantly different between donors for IP, oRG, and CPN cells (**Fig. 5C**, **Suppl. Table 3**).

Finally, we examined whether there were donor-specific differences in cell-intrinsic expression changes in the VPA-treated Chimeroids. To summarize the magnitude of differential response in each donor, we calculated pseudobulk expression profiles for each donor within each Chimeroid, both globally (summing the UMI counts of all cells from a donor in a Chimeroid) and separately for each major cell subset (e.g., summing the UMI counts of aRGs from one donor in a Chimeroid). For each set of pseudobulk expression profiles (global, aRG, oRG, IP, CFuPN, and PN), we performed principal component analysis (PCA) to identify the major axes of variation, and then compared the Euclidian distance between control and VPA-treated Chimeroids for each donor, calculated on all principal components (**Methods**). The CW and Mito210 donors showed less separation between VPA and control than did the other donors in global profiles (**Fig. 5D**). To examine the impact on intrinsic cell states specifically, we performed this analysis for each affected cell subset separately showing that donors also differed in the degree of expression response to VPA within individual cell types (**Fig. 5D**). For most cell types, the 11a donor was among the most-affected, while CW was among the least-affected; however, for CFuPNs, CW showed the second-strongest effect, indicating that individual cell types within each donor show different degrees of perturbation, a finding consistent with our prior report that the effect of genetic background is cell-type specific and perturbation-specific (Paulsen et al., 2022).

Altogether, our data show that the multi-donor Chimeroid system allows assaying experimental perturbations across multiple donors while still enabling the detection of individual differences in response. The data highlight the value of Chimeroids for high-throughput investigation of the effect of human genetic variation on brain responses.

## Discussion

Human genetic diversity is an important contributor to clinical variation in disease manifestation, but modeling the contribution of genetic background to disease phenotypes of the human brain has been difficult due to the requirement for human experimental brain models that can also accommodate cells from a large variety of individual donors. Not all human PSC lines perform equally well across all *in vitro* models, especially organoid systems, due to differences inherent to each line (Wells et al., 2023), such as biases in fate potential and proliferation linked to differential reprogramming, epigenetic imprinting, or susceptibility to culture conditions.

Our demonstration that cells derived from multiple donors can grow together and are capable of balanced differentiation within individual Chimeroids indicates that it is possible to investigate, in a single organoid, the collective development and behavior of a large diversity of cell types of the brain, across many human individuals. Because Chimeroid protocols can be adapted to tolerate even extreme donor-to-donor differences, while still allowing detection of differential response to perturbation, donors can be mixed agnostically to stem cell line of origin. This paves the way for efforts over the coming years aimed at scaling-up the donor pools used in Chimeroids to enable investigation of cells from a very large numbers of individuals.

The Chimeroid approach will facilitate assaying across a variety of donor characteristics; for instance, panels of lines that vary in genetic risk for disease, presence or absence of predisposing factors, or clinical phenotype. Combining highly-multiplexed Chimeroids with increased automation of cell culture systems may enable sufficient scaling for investigation of population-level variation across hundreds or even thousands of individuals.

Our demonstration that variation in donor response to perturbations such as neurotoxic compounds can be read at cell-type resolution, suggests that Chimeroids may be also used to stratify patients across treatment response groups. Therapeutics for CNS diseases often perform poorly in clinical trials and typically show efficacy in only a subset of patients. The ability to assay drug response at scale using Chimeroids may enable data-driven stratification of patients into different response groups, and in time would produce matrices of drug response data by cell type and by donors to fuel generative models that may predict drug efficacy before clinical investigation.

## Acknowledgements

We thank J. R. Brown (from the P.A. laboratory) for input and assistance in editing the manuscript and all the members of the Arlotta laboratory for discussions; B. Cohen (McLean Hospital) for the Mito210 iPS line; G. Church (Harvard University) for the PGP1 iPS line; M. Talkowski (MGH) for the GM08330 iPS line; the Broad Genomics Platform for sequencing; and Steve McCarroll (Harvard Medical School) for the Census-seq protocol. This work was supported by grants from the Stanley Center for Psychiatric Research to P.A., R.N., and J.Z.L., the Broad Institute of MIT and Harvard, the National Institutes of Health (P50-MH094271 and RF1-MH123977 to P.A., R01-MH112940 to P.A. and J.Z.L., and U01-MH115727 to R.N.), and the Klarman Cell Observatory (A.R.).

## Authors’ contribution

N.A.B, I.F and P.A. conceived and designed the experiments. N.A.B, I.F and, S.A generated, cultured, and characterized all the organoids in this study, with the help of S.T. and R.K., and P.A. supervised their work and contributed to data interpretation. X.A. and D.J.D.B. contributed to some of the scRNA-seq experiments, with help from N.A.B and I.F and supervision of P.A. and J.Z.L. N.A.B., I.F and S.T. performed scRNA-seq and library preparation for the vast majority of the preparations included in this study. T.F., N.A.B and I.F worked on cell type assignments and scRNA-seq data analysis, and T.F performed all the computational work under supervision by A.R and P.A.; R.N. and M.T. provided the CW line, and advised on the PSC-Chimeroid experiments and on application of the Census-seq analysis pipeline, using computational tools and resources provided by Steven McCarroll’s laboratory. N.A.B, I.F. and P.A. wrote the manuscript with contributions from all authors. All authors read and approved the final manuscript.

## Materials & Correspondence

Correspondence and request for materials should be addressed to P.A. Reprints and permissions information is available at www.nature.com/reprints.

## Code Availability

Code used during data analysis is available at [available before publication].

## Data Availability

Read-level data from scRNA-seq have been deposited in a controlled access repository at [available before publication] while count-level data and meta-data have been deposited at the [available before publication]. Any additional information required to reanalyze the data reported in this paper is available from the lead contact upon request. Data from previous publications that were used in this study can be found at [available before publication].

## Methods

### Human pluripotent stem cell culture

All human pluripotent stem cell lines were cultured as previously described (Paulsen et al., 2022; Uzquiano et al., 2022; Velasco et al., 2019). Briefly, MTESR1 medium (StemCell Technologies), mTESR+ medium (StemCell Technologies), or StemFlex medium (Gibco), all with 1% of added Penicillin-Streptomycin Solution (Corning), were used for culture of stem cells in cell culture dishes (Falcon) previously coated with 1% Geltrex (Gibco), at 37°C in 5% CO2. All hPS cell lines were maintained below passage 50, tested negative for mycoplasma (assayed with MycoAlert PLUS Mycoplasma Detection Kit, Lonza) and were karyotypically normal (G-banded karyotype test performed by WiCell Research Institute).

### Characterization of the PSC lines

The psychiatric control Mito210 male iPS cell line was provided by B. Cohen (McLean Hospital); the PGP1 male iPS cell line (Church, 2005) was provided by G. Church (Harvard University); the GM08330 male iPS cell line (Sheridan et al., 2011) was provided by M. Talkowski (MGH) and was originally derived from fibroblasts obtained from the Coriell Institute for Medical Research (Sugathan et al., 2014). The female CW50037 iPSC line was from the California Institute for Regenerative Medicine (CIRM) iPSC collection. The H1 male hESC line (also known as WA01) (Thomson et al., 1998) was purchased from WiCell; and the 11a male iPS cell line(Boulting et al., 2011) was obtained from the Harvard Stem Cell Institute.

All lines were authenticated as follows: The PGP1 iPSC line (performed by TRIPath in 2018) (Uzquiano et al., 2022). The Mito210 iPSC line was authenticated with genotyping analysis (Fluidigm FPV5 chip) performed by the Broad Institute Genomics Platform (Uzquiano et al., 2022). The H1 and GM08330 lines were authenticated by STR analysis (performed by WiCell in 2021). The GM08330 parental line has a previously-reported interstitial duplication in the long (q) arm of chromosome 20 (Paulsen et al., 2022); all other lines were karyotypically normal. All experiments involving human cells were approved by the Harvard University IRB and ESCRO committees.

### Cortical organoid differentiation

Dorsally patterned forebrain organoids were generated according to (Velasco et al., 2019), with some adaptions to the protocol as previously described (Paulsen et al., 2022; Uzquiano et al., 2022). Briefly, on day 0, feeder-free cultured human iPSCs or hESCs, 75–85% confluent, were enzymatically dissociated to single cells with Accutase (Gibco), and 9,000 cells per well were reaggregated in ultra-low cell-adhesion 96-well plates with V-bottomed conical wells (sBio PrimeSurface plate; Sumitomo Bakelite) in the same pluripotent cells medium in which they were previously maintained. At day 1, 80 μl of media was replaced with Cortical Differentiation Medium (CDM) I, containing Glasgow-MEM (Gibco), 20% Knockout Serum Replacement (Gibco), 0.1 mM Minimum Essential Medium non-essential amino acids (MEM-NEAA) (Gibco), 1 mM pyruvate (Gibco), 0.1 mM 2-mercaptoethanol (Gibco), with 1% of Penicillin-Streptomycin Solution (Corning). From day 0 to day 6, ROCK inhibitor Y-27632 (Millipore) was added to the medium at a final concentration of 20 μM. Patterning small molecules, Wnt inhibitor IWR1 (Calbiochem) and TGFβ inhibitor SB431542 (Stem Cell Technologies), were added from day 0 to day 15-18, at a concentration of 3 μM and 5 μM, respectively.

From day 15-18, patterned EBs were cultured under orbital agitation (Thermofisher CO2 resistant orbital shaker) in ultra-low attachment culture dishes (Corning) in CDM II medium, containing DMEM/F12 medium (Gibco), 2mM Glutamax (Gibco), 1% N2 (Gibco), 1% Chemically Defined Lipid Concentrate (Gibco), 0.25 μg/mL fungizone (Gibco), with 1% of Penicillin-Streptomycin Solution (Corning). On day 35, cell aggregates were transferred to spinner-flask bioreactors (Corning) and maintained in CDM III (CDM II supplemented with 10% fetal bovine serum (FBS) (GE-Healthcare), 5 μg/mL heparin (Sigma) and 1% Matrigel (Corning)). From day 70, organoids were cultured in CDM IV (CDM III supplemented with B27 supplement (Gibco) and 2% Matrigel).

### Multi-donor cortical Chimeroids generation

Chimeroid generation is based on a reaggregation process using putative NSCs or NPCs originating from dorsally-patterned forebrain organoids (cortical organoids) dissociated at distinct developmental stages. Pools of cortical organoids from each donor were enzymatically dissociated into a single-cell suspension using the Worthington Papain Dissociation System kit (Worthington Biochemical). Cell suspensions derived from different donors were mixed at equal ratios, and 18,000-20,000 cells per well were reaggregated in ultra-low cell-adhesion 96-well plates with V-bottomed conical wells (sBio PrimeSurface plate; Sumitomo Bakelite). The following day, the media was replaced with 100 μl of CDM2 per well. Two days after mixing, reaggregated EB’s were passaged in ultra-low attachment culture dishes (Corning), and cultured under orbital agitation (Thermofisher CO2 resistant orbital shaker). The subsequent media changes of the Chimeroids followed our published protocol changing to CDM3 media at DIV35, and then to CDM4 at DIV70.

### EtOH and VPA treatment

At DIV45, multidonor and single-donor Chimeroids were treated with either EtOH or VPA. EtOH (E7023-6, Millipore-Sigma) was administered directly to the culture media every two days at a concentration of 50 μM and the plate was sealed with Parafilm M (Bemis) for 24 hours after administration. VPA (P4543-25G, Millipore-Sigma) was added to the culture media at a concentration of 0.7 mM and the media was changed twice per week. Both treatments were stopped at DIV75, and organoids were analyzed at 3 mo.

### Fixation and processing of samples for cryosectioning

Chimeroids were fixed in 4% paraformaldehyde (PFA) (Electron Microscopy Services) overnight (O/N) in a 12-well plate (Falcon) at 4°C, washed with 1X phosphate buffered saline (PBS) (Gibco) 3 times, and cryoprotected in a 30% sucrose solution (Sigma) in PBS (O/N) at 4°C. Samples were then embedded in bovine gelatin. Gelatin solution containing 10% of bovine gelatin (Sigma) and 7.5 % of sucrose (Sigma) was prewarmed at 37°C for 15 min. The 30% sucrose solution was removed from the samples and exchanged for the prewarmed gelatin, and the samples were incubated at 37°C for 15 min. Meanwhile, plastic molds were coated with a 2mm layer of warm gelatin solution and left to polymerize at room temperature (RT). Chimeroids were transferred to the pretreated plastic molds, and 1 mL of warm gelatin solution was added to cover the Chimeroids. Following polymerization at RT for 3 min, Chimeroids were pre-chilled at 4°C for 15-20 min. Finally, the molds were frozen in a cold bath containing 100% EtOH and dry ice for 2-3 min, and stored at -80°C indefinitely.

### Immunohistochemistry

For immunohistochemistry, 14 to 20 μm thick sections were cut using a cryostat (Leica). Cryosections were stabilized at RT for 5 min and blocked with 10% donkey serum (Sigma) + 0.3% Triton X-100 (Sigma) in PBS. Primary antibodies (**Suppl. Table 4**) were diluted in the blocking solution and incubated O/N. After 4 washes with PBS, cryosections were incubated at RT with secondary antibodies diluted in PBS (1:1000; **Suppl. Table 4**) for 1 hour at room temperature, washed 4x with PBS, and stained with DAPI (1:10,000 in PBS + 0.1% Tween-20) for 15 minutes to visualize cell nuclei.

### Microscopy

Immunofluorescence images were acquired with a Zeiss Axio Imager.Z2 and Lionheart™ FX Automated Microscope (BioTek Instruments). Images were analyzed and processed with the Gen5 (BioTek Instruments) and Zen Blue (Zeiss) image processing software, followed by ImageJ (Fiji)(Schindelin et al., 2012).

### Dissociation of brain organoids and scRNA-seq

Organoids were dissociated as previously described (Velasco et al., 2019), with the exception of the 1-mo organoids, which were dissociated using low-binding 1.5 mL tubes (Eppendorf). Dissociated cells were resuspended in 100 μl of PBS, filtered with a 35 μm cell strainer tube (Corning) to avoid aggregates, and counted with a Cellometer K2 instrument (Nexcelom) or a Countess II automated hematocytometer (Thermo Fisher Scientific). Single-cell suspensions were loaded into the Chromium Next GEM Chip G (10x Genomics, 1000120) and run with the Chromium Controller to generate single-cell GEMs. scRNA-seq libraries were prepared with the Chromium Single Cell 3’ Library and Gel Bead Kit v3.1 (10x Genomics, 1000268). When Chimeroids are composed of cells from multiple genetic backgrounds, it is possible to use that genetic variation to identify doublets containing cells from two different cell lines. This enabled us to tolerate a higher doublet rate and load ∼20,000-30,000 cells per channel. The resulting libraries were pooled based on molar concentration and sequenced on a NextSeq 500 or NovaSeq 6000 instrument (Illumina) with 28 bases for read 1, 55 bases for read 2 and 8 bases for index read 1. If necessary, after the first round of sequencing, we re-pooled libraries based on the actual number of cells in each library, and re-sequenced with the goal of producing an equal number of reads per cell for each sample (with a target read depth of 20,000 reads/cell).

### Single Cell RNA-Seq data processing

All scRNA-seq data were processed using 10x Genomics Cell Ranger v7.1.0(Zheng et al., 2017). Raw BCL files were converted to FASTQ files using the *mkfastq* command with default parameters. Cellranger’s *count* function was used on the resulting FASTQ files to produce cell-by-gene count matrices for each organoid, with the pre-built GRCh38 human reference genome provided on the 10x Genomics downloads page (https://support.10xgenomics.com/single-cell-gene-expression/software/downloads/latest GENCODE v32/Ensembl 98). The *expect-cells* flag was set manually for each sample, with values ranging from 10,000 to 30,000. All other parameters were set to default. Cellrangers’s filtered_feature_bc_matrix was imported into Seurat v4.3.0 (Stuart et al., 2019) using R version 4.2.2 for downstream analyses.

### Genetic demultiplexing

For genetic demultiplexing based on scRNA-seq data, demuxlet was applied (Kang et al., 2018), with default settings, to the alignments produced by Cellranger. Reference variant-call-format (VCF) files generated from whole-genome sequencing of each donor cell line were processed with bcftools (Danecek et al., 2021) to normalize the reference VCFs into biallelic variants, and remove non-passing and monomorphic variants. Droplets that demuxlet determined to have ambiguous genetic origins, or which were likely to be heterogeneous doublets, were removed with default settings from the Seurat objects before downstream analysis.

In order to determine the proportional abundance of each donor in PSC, NSC, and NPC Chimeroids at different timepoints and from different mixes, Census-seq was applied to low-pass WGS data (Wells et al., 2023), using the Census-seq function from the Drop-seq software package with default parameters, following the Census-seq computational protocols found at https://github.com/broadinstitute/Drop-seq, and using the same reference VCF files as mentioned above for each donor.

### Cell profile quality filtering, normalization, clustering, annotation, and integration

For each Chimeroid, cell profiles with fewer than 200 expressed genes, fewer than 500 UMIs, more than 20,000 UMIs, or greater than 15% mitochondrial RNA, were removed. Seurat’s *SCTransform* function was then run on each organoid’s filtered counts matrix, regressing out the percent mitochondrial RNA. Principal component analysis (PCA) was performed on the variable features found by SCTransform. Seurat’s *FindNeighbors* function was used to construct a *k*-nearest-neighbor graph (*k*=20) with the top 30 PCs, followed by Louvain clustering using the *FindClusters* function, with resolution set between 0.2 and 1.0.

Single-cell expression data from multiple Chimeroids were combined into several larger joint datasets based on protocol and treatment as follows: PSC-Chimeroids of Mix 1, Mix 3, and Mix 4 (n=5; 62,242 cells); PSC-Chimeroids of Mix 5 (n=2; 15,612 cells); control, EtOH-treated, and VPA-treated NSC-Chimeroids of Mix 1 and Mix 2 (n=21; 200,019 cells); NSC- and NPC-Chimeroids of Mix 3 and Mix 4 (n=6; 52,771 cells); and control, EtOH-treated, and VPA-treated single-donor NSC-Chimeroids (n=45; 112,166 cells).

To visualize scRNA-seq data from multiple organoids in shared UMAP embeddings, raw UMI counts matrixes were merged using Seurat’s *merge* function, then jointly renormalized with SCTransform. PCA was performed on the joint expression matrix, followed by *k*-nearest-neighbor graph construction and clustering using the same methods as on the individual datasets. When merging all control, VPA-treated, and EtOH-treated Mix 1 and Mix 2 NSC Chimeroids, there was a clear batch effect which resulted in two of the organoids clustering entirely separately from all other organoids. In this case, dataset integration was performed as in Seurat’s integration vignette using SCTransform; specifically, using the functions *SelectIntegrationFeatures* (with the *nfeatures* parameter set to 3000), *PrepSCTIntegration*, *FindIntegrationAnchors*, and *IntegrateData*. PCA was performed on the integrated dataset, and cell profiles (top 30 PCs) were embedded by Uniform Manifold Approximation and Projection (UMAP). Within each dataset, each cluster was manually annotated with a cell-type label based on a combination of (1) canonical marker gene expression, (2) examining a list of upregulated genes within each cluster as determined by Seurat’s *FindMarkers* function, and (3) automated label transfer from the expression profiles of our previously-published organoid atlas (Uzquiano et al., 2022) using Seurat’s *FindTransferAnchors* and *MapQuery* functions. Cell-type labels were further refined in each merged or integrated dataset based on the joint cluster labels. Cell profiles (top 30 PCs) were embedded by UMAP. Clusters that had exceptionally low UMI counts were labeled as low-quality cells and removed from downstream analysis. Clusters that initially could only be described as cycling progenitors were subclustered by subsetting to only the cells from those clusters and rerunning PCA, *k*-nearest-neighbors graph construction, and clustering; expression of HOPX, EOMES, and SOX2 was used to identify these subclusters as aRG, oRG, or IP. In the case of the single-donor NSC-Chimeroids, one cluster which included both cells that expressed DLX2 and cells that expressed RELN was subclustered to distinguish immature interneurons and Cajal-Retzius cells.

### Pseudotime analysis

Monocle3 version 1.1.3 (Cao et al., 2019; Qiu et al., 2017; Trapnell et al., 2014) was applied to the scRNA-seq data from 3-mo, Mix 3 NSC-Chimeroids, using aRGs as the root cells.

### Reproducibility and comparison with fetal tissue and cortical organoids

Adjusted Mutual Information (AMI) scores were calculated between donor source and cell type calls for the cells within each batch of multi-donor Chimeroids and single-donor Chimeroids, using the *aricode* R package version 1.0.2(*CRAN - Package Aricode*, n.d.). Seurat’s *FindTransferAnchors* and *MapQuery* functions were used as described in the Seurat 4.3 label-transfer tutorial to transfer labels from fetal cortex samples and cortical organoids onto Chimeroids and single-donor Chimeroids (and vice-versa).

Seurat’s *FindMarkers* function, with no fold-change or significance threshold, was used to independently create a list of marker genes for each cell type in untreated NSC-Chimeroids, fetal cortical tissue or cortical organoids, ranked by significance as calculated via Wilcoxon rank-sum test, with the most significantly upregulated genes within a cell type at the top of the list, and the most significantly downregulated genes at the bottom. The improved rank-rank hypergeometric overlap test from the RRHO2 R package (version 1.0) was used to compare the marker gene lists from each NSC-Chimeroid cell type to those for each fetal and organoid marker list.

For each of 19 canonical cell type marker genes, the mean scaled expression across cells of each of 5 major cell types (aRG, oRG, IP, CFuPN, and CPN) was calculated in each of 5 datasets (Mix 1/Mix 2 NSC-Chimeroids, NPC-Chimeroids, single-donor Chimeroids, atlas organoids, and endogenous fetal cortex), in order to compare marker gene expression between datasets within comparable cell types. Pearson’s correlation coefficient of those expression values was calculated between the endogenous fetal tissue and the corresponding cell types in each organoid and Chimeroid dataset, and also between the atlas organoids and each Chimeroid dataset.

### Cell type abundance, donor abundance, and neighborhood changes between treatment conditions

A negative binomial mixed-effects model was used to test for differential cell type abundances between control Chimeroids and VPA or EtOH-treated Chimeroids. Specifically, the *glmer.nb* function from lme4 (version 1.1.33) was applied to a matrix of cell type counts per donor within each Chimeroid, using the log of the total number of cells per donor and Chimeroid as an offset, Chimeroid differentiation batch and donor Mix (Mix 1 or Mix 2) as fixed effects, and both donor identity (to account for general differences in cell type abundances between donors and similarities within donors across replicates) and individual Chimeroid (to account for similarities in cell type abundance between donors within a shared organoid) as random effects. Thus, a model formula of Cell Type ∼ Treatment + offset(log(LibrarySize)) + Batch + Mix + (1|Chimeroid) + (1|Donor) was used. Significance of the effect of treatment on cell type abundance was calculated by comparing this model with a model without the fixed effect for treatment, but which was otherwise identical, via the *anova* function in the R *stats* package, version 4.2.2. Benjamini-Hochberg (BH) False Discovery Rates (FDR) were calculated based on the resulting p-values.

Additional negative binomial mixed-effects models were used to test whether the magnitude and/or direction of change in cell type in response to VPA or EtOH treatments were different between the different donors. Specifically, models similar to those described above were implemented, but with an additional random slope parameter allowing for the treatment effect to vary by donor. Thus, the *anova* function was used to compare a model with the formula: Cell Type ∼ Treatment + offset(log(LibrarySize)) + Batch + Mix + (1|Chimeroid) + (Treatment|Genotype) to an otherwise identical model where the (Treatment|Genotype) term was replaced with (1|Genotype). In cell types where the inclusion of the random slope term significantly improved the model (oRGs, IPs, and CPNs), post-hoc pairwise tests were performed between each pair of donors using the same mixed-effects models.

EdgeR version 3.40.2 (Chen et al., 2016; McCarthy et al., 2012; Robinson et al., 2010) was used to test for changes in donor abundances in response to VPA or EtOH treatments. Specifically, a matrix of cell counts per donor within each Chimeroid was passed through the functions calcNormFactors (with method set to “TMM”), estimateDisp (with trend set to “none”), glmQLFit (with abundance.trend set to “FALSE”), and glmQLFTest, using a design formula with Chimeroid differentiation batch and donor Mix as additional covariates. Significance was reported as FDR, as calculated via BH.

The miloR R package version 1.6.0 (Dann et al., 2022) was used to assess differential abundance within overlapping cell neighborhoods, ranging in size from 50-200 cells each. For each treatment (VPA and EtOH), the expression matrix was subset to include only the top 3,000 most variable genes within cells from control and treated Chimeroids, PCA was performed on that submatrix, and the result was passed into a Milo object for further analysis. miloR’s *buildGraph* function was run with *k*=30 and *d*=30, followed by *makeNhoods* and *calcNhoodDistance*. Differential abundance (DA) testing was performed for treated *vs*. control cells within each neighborhood using *testNhoods* with a design of ∼Mix + Donor + Treatment. Significance is reported as a spatial FDR value calculated by miloR based on DA p-values and relationships between neighborhoods.

### Pseudobulk profiles and differential expression analysis

Pseudobulk profiles were generated both globally (wherein a pseudobulk profile comprises the UMIs from all cells from one donor within one chimeroid), and on a cell type-by-cell type basis (wherein a pseudobulk profile comprises the UMIs from all cells of one type from one donor within one chimeroid), by summing UMI counts for each gene and, where appropriate, calculating expression values as the log-Counts-Per-Million within each pseudobulk profile. Pseudobulk profiles comprising fewer than 20 cells were removed from downstream analyses.

Principal Component Analysis (PCA) was performed on highly variable genes (defined as genes where the mean expression plus the log of the variance-to-mean expression ratio is greater than 1) of the global pseudobulk expression matrices for the global and each cell type expression profiles. For each analysis, outliers (2-8 per expression matrix) were identified and removed by calculating the Euclidian distance between each sample and the centroid of all samples; outliers were defined as samples where this distance was more than 2.5 times the interquartile range above the upper quartile bound. PCA was re-run on the remaining samples for downstream analysis. The Euclidean distances between each pseudobulk profile across all PCs (the number of PCs in each case equaled the number of pseudobulked samples which passed quality and outlier filters) within each PCA were calculated using the dist() function from the *stats* R package. Analysis of Variance (ANOVA) was performed to assess whether distances between control and treated samples were significantly different between the five donor lines. When the ANOVA yielded a p-value of less than 0.05, post-hoc Tukey tests were performed for pairwise comparisons between the donors.

## Figure Legends

**Suppl. Fig. 1.**
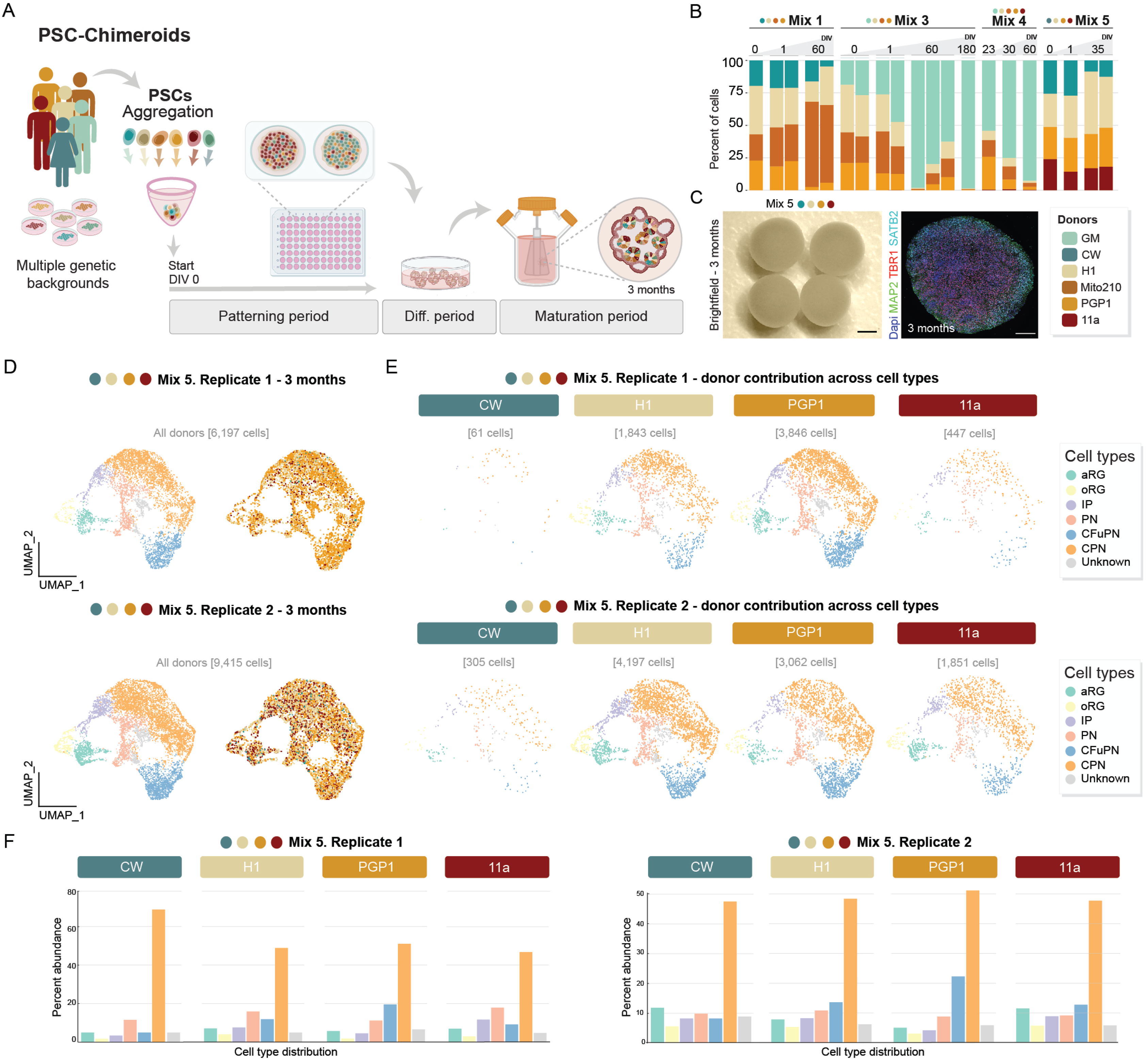
PSC Chimeroids display uneven donor composition but proper cell type composition. **A,** Schematic of the PSC-Chimeroid protocol. **B**, Stacked bar plots showing donor composition, as measured via Census-seq from low-pass whole-genome sequencing, in PSC-Chimeroids from various mixes and timepoints, labeled as the number of days since the day of the seeding (DIV0). For the DIV0 and DIV1 time points, n=4 chimeroids have been pulled together. **C**, Left panel: brightfield image of 3-mo PSC-Chimeroids; scale bar: 1 mm. Right panel: immunolabelling of 3-mo PSC-Chimeroid with SATB2, TBR1 and MAP2. Scale bar: 500 μm. **D**, UMAP of integrated PSC-Chimeroids at 3 mo, colored by annotated cell type (left panel) and donor line as determined with demuxlet (right panel). **E**, UMAPs split by donor for two different replicates. **F**, Barplots of cell type proportions in PSC Chimeroids (n=2), demultiplexed by donor. DIV, days *in vitro*.

**Suppl. Fig. 2.**
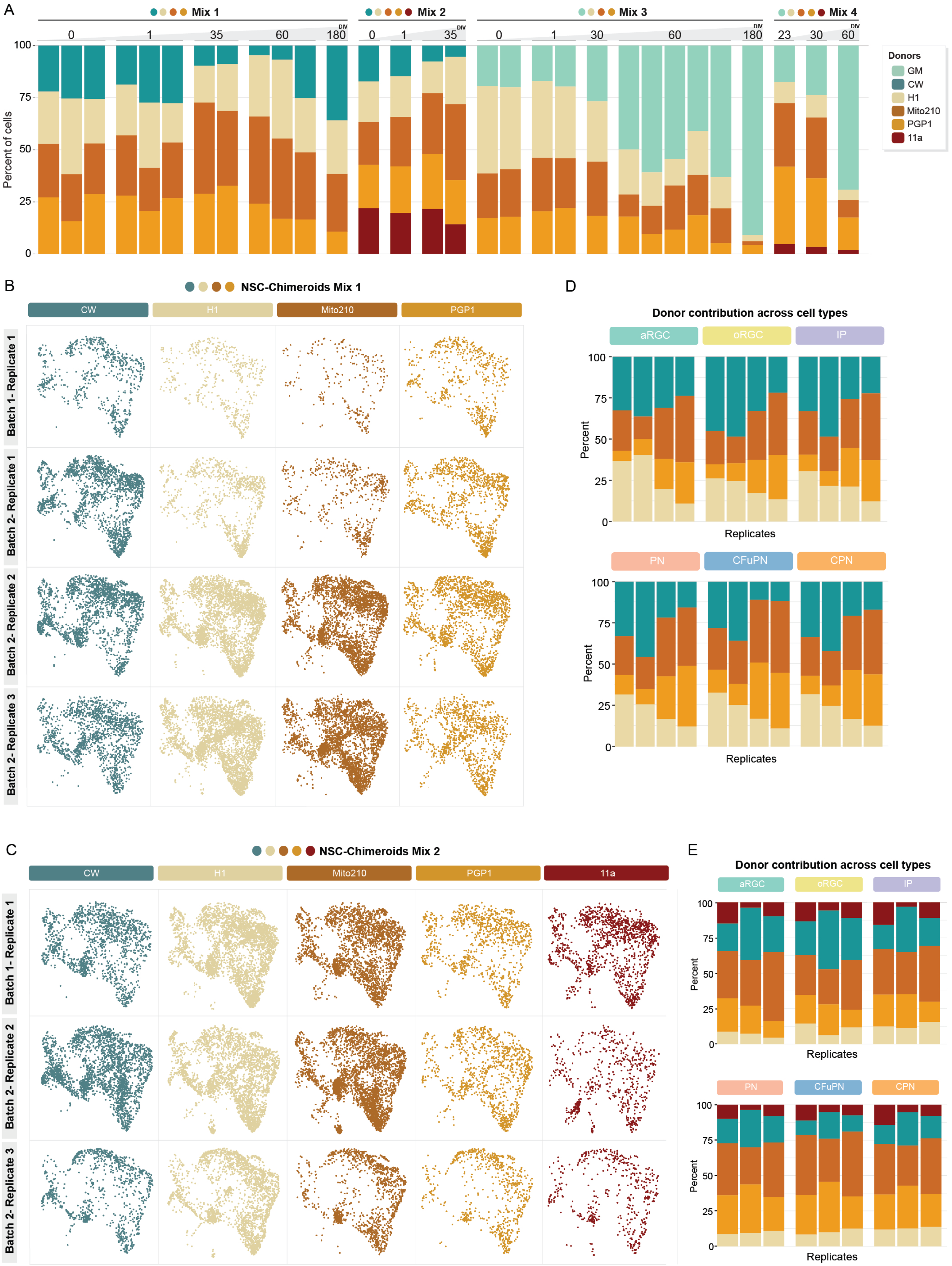
NSC-Chimeroids develop proper cell-type composition across multiple donors. **A**, Stacked barplots showing the donor composition, as measured via Census-seq from low-pass whole-genome sequencing, of multi-donor NSC-Chimeroids from various mixes and at various timepoints, labeled as the number of days since reaggregation (DIV; for the DIV0 and DIV1 time points, n=4 chimeroids have been pulled together). **B-E,** 3-mo NSC-Chimeroids UMAPs split by donor (**B, D**) and stacked barplots of donor contributions for each cell type (**c, e**) for Mix 1 (4 donors) and Mix 2 (5 donors).

**Suppl. Fig. 3.**
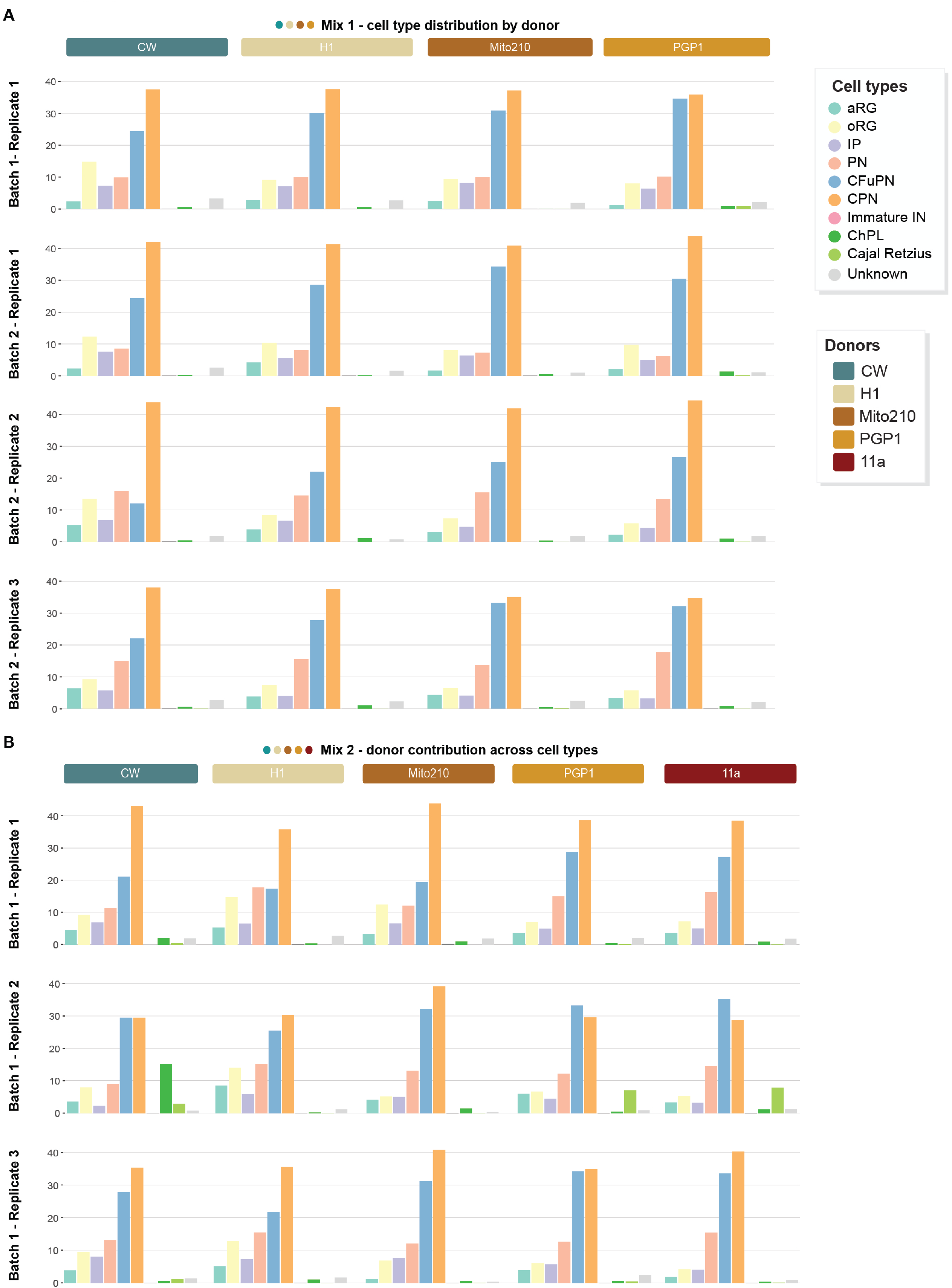
Individual donors within NSC-Chimeroids maintain consistent proportions of cortical cell types. **A-B**, Distribution of cell types across individual donors, across two independent mixes, Mix 1 (**A**) and Mix 2 at 3 mo (**B**). Donors were demultiplexed using demuxlet.

**Suppl. Fig. 4.**
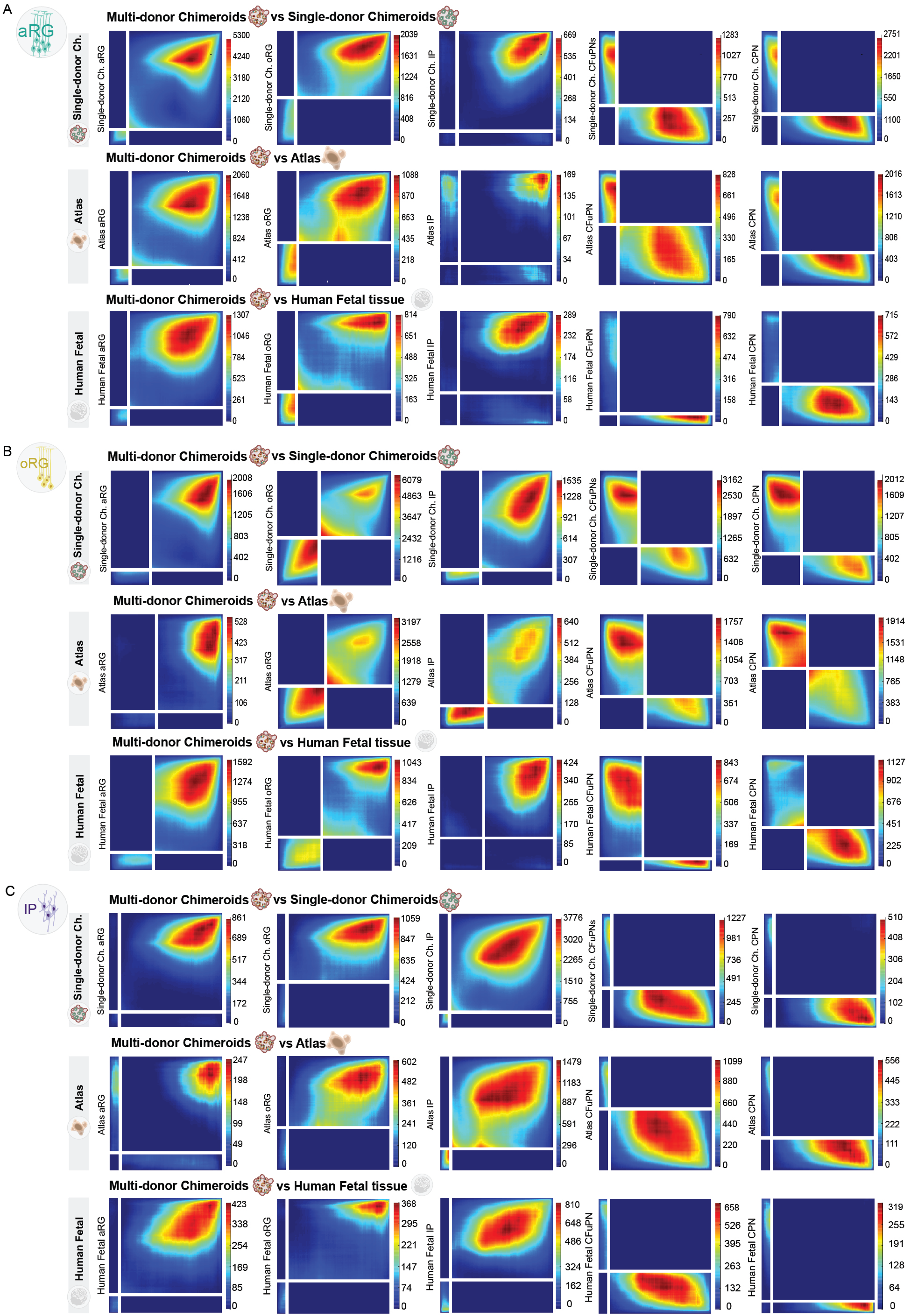
Chimeroids display similar neural progenitor complexity as single-donor organoids and fetal tissue. Rank-Rank Hypergeometric Overlap (RR-HO) plots comparing the expression signatures of progenitor types in multi-donor Chimeroids to all cell types in single-donor chimeroids, atlas cortical organoids or endogenous human fetal tissue. The horizontal axes represent lists of marker genes for multi-donor Chimeroid progenitor types (**A-C**; aRG, oRG and IP) compared to all other Chimeroid cells, ranked from most upregulated to most downregulated; the vertical axes represent similarly ranked lists of marker genes for, single-donor Chimeroids, atlas organoids or human fetal cells. Color at a given position represents the significance (negative log p-value) of the overlap of the gene lists up to that point, as calculated by Fisher’s exact tests. High significance (i.e., red color) in the lower left and upper right quadrants indicates strong concordance between the expression profiles which define the compared cell types.

**Suppl. Fig. 5.**
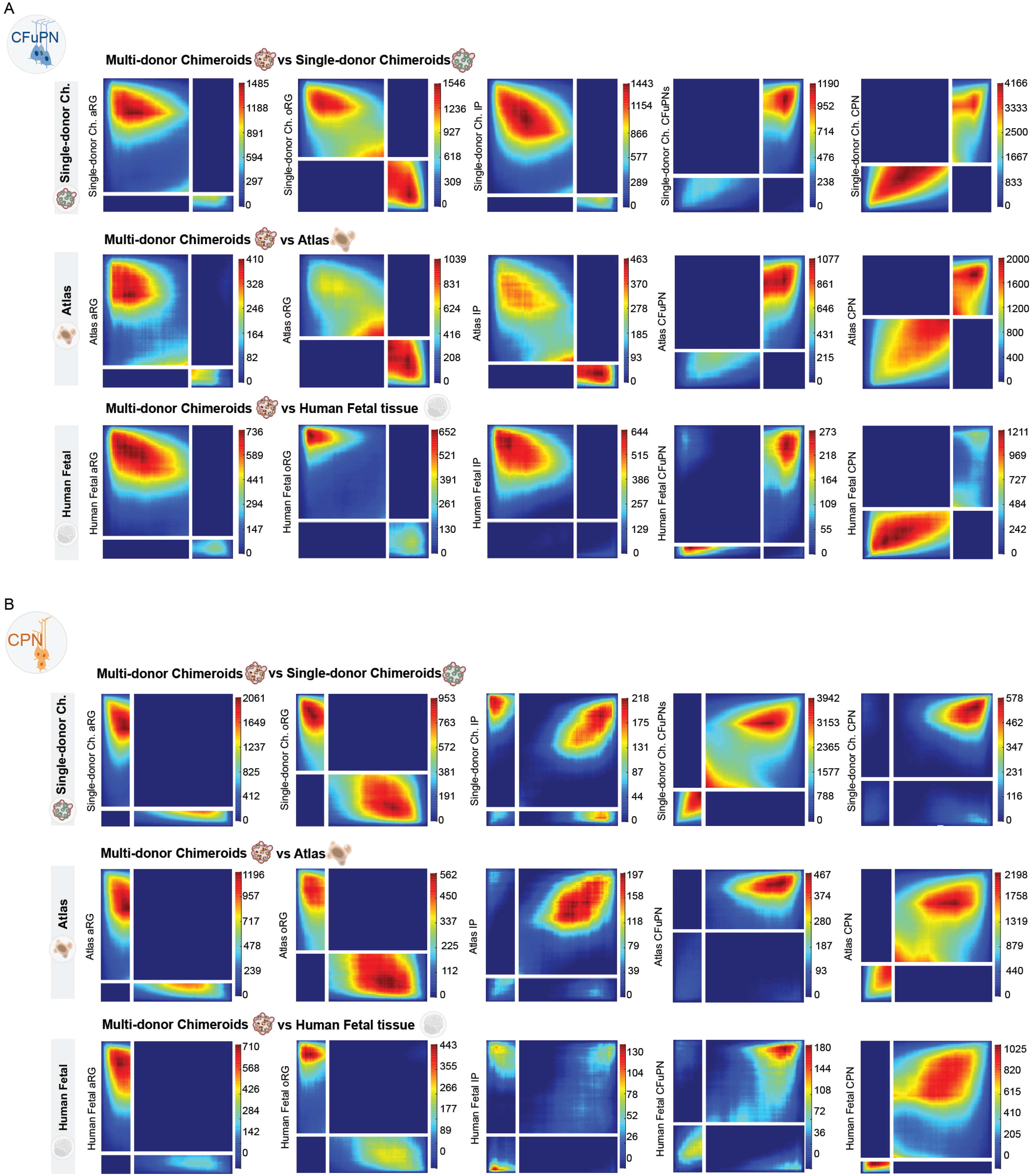
Chimeroids display similar neuronal complexity as single-donor organoids and fetal tissue. RR-HO plots comparing the expression signatures of pyramidal neuron types in multi-donor Chimeroids to all cell types in single-donor Chimeroids, atlas cortical organoids or endogenous human fetal tissue. The horizontal axes represent lists of marker genes for multi-donor Chimeroid pyramidal neuron types (**A,B**; CFuPN and CPN) compared to all other Chimeroid cells, ranked from most upregulated to most downregulated; the vertical axes represent similarly ranked lists of marker genes for, single-donor Chimeroids, atlas organoids or human fetal cells.

**Suppl. Fig. 6.**
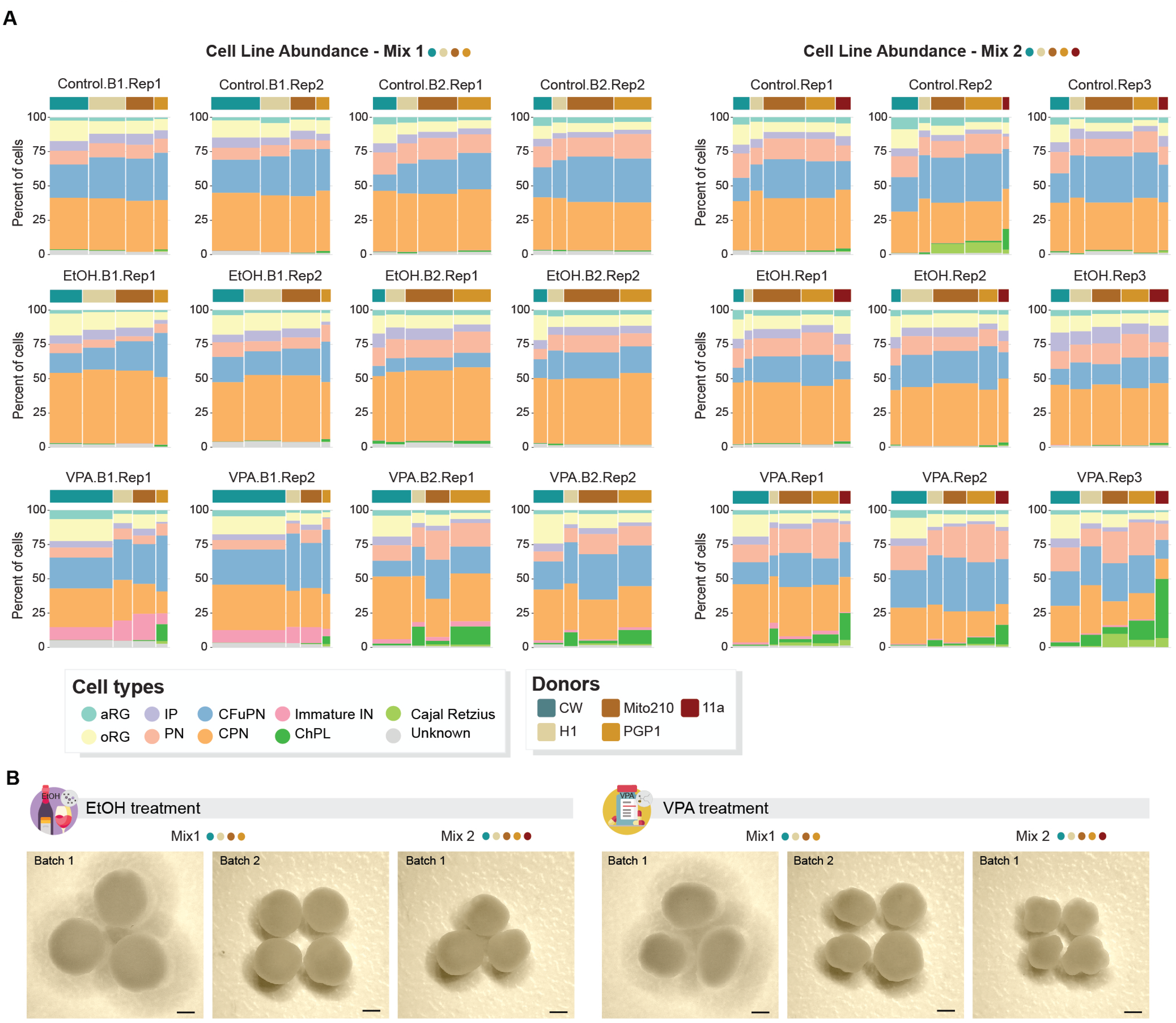
VPA and EtOH treatment differently affect cell-type composition in NSC-Chimeroids. **A**, Stacked barplot showing cell type and donor composition for treated and control Chimeroids; the width of each bar corresponds to the proportion of the indicated donor in Mix 1 (4 donors, left panel) and Mix 2 (5 donors, right panel). **B**, Brightfield images of whole cortical NSC-Chimeroids at 3 mo, in the EtOH and VPA treatment conditions. Scale bar: 1 mm.

**Suppl. Fig. 7.**
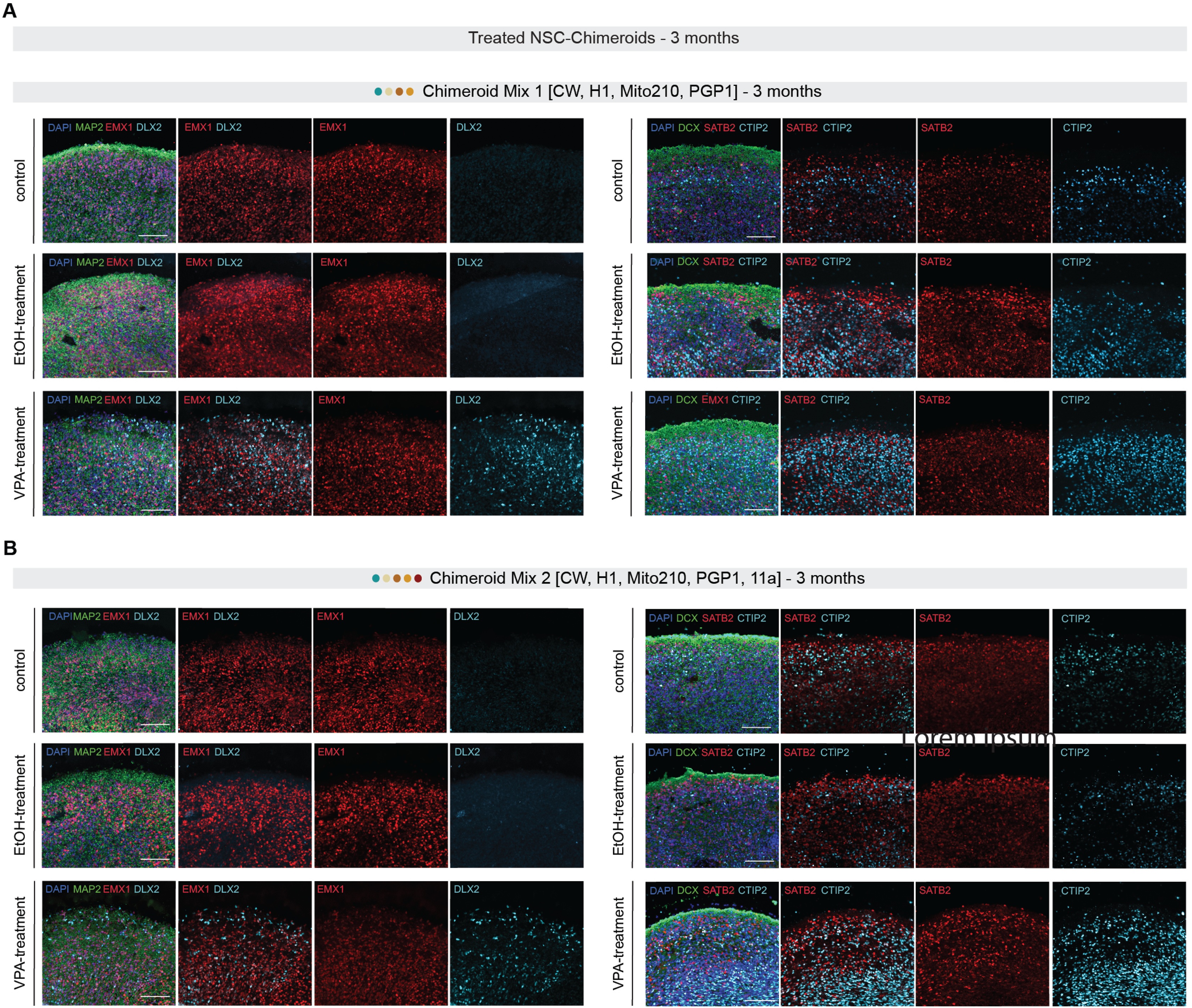
VPA-treated Chimeroids develop an immature interneuron population. Immunolabelling of multi donor NSC-Chimeroids (**A**, Mix 1, upper images and **B**, Mix 2, lower images). Lefthand panel, MAP2 (neuronal dendrites), EMX1 (cortical progenitors), and DLX2 (GABAergic cells). Righthand panel, DCX (migrating neurons), SATB2 (upper layers cortical neurons), and CTIP2 (deep layers cortical neurons). Scale bar: 100 μm.

**Suppl. Fig. 8.**
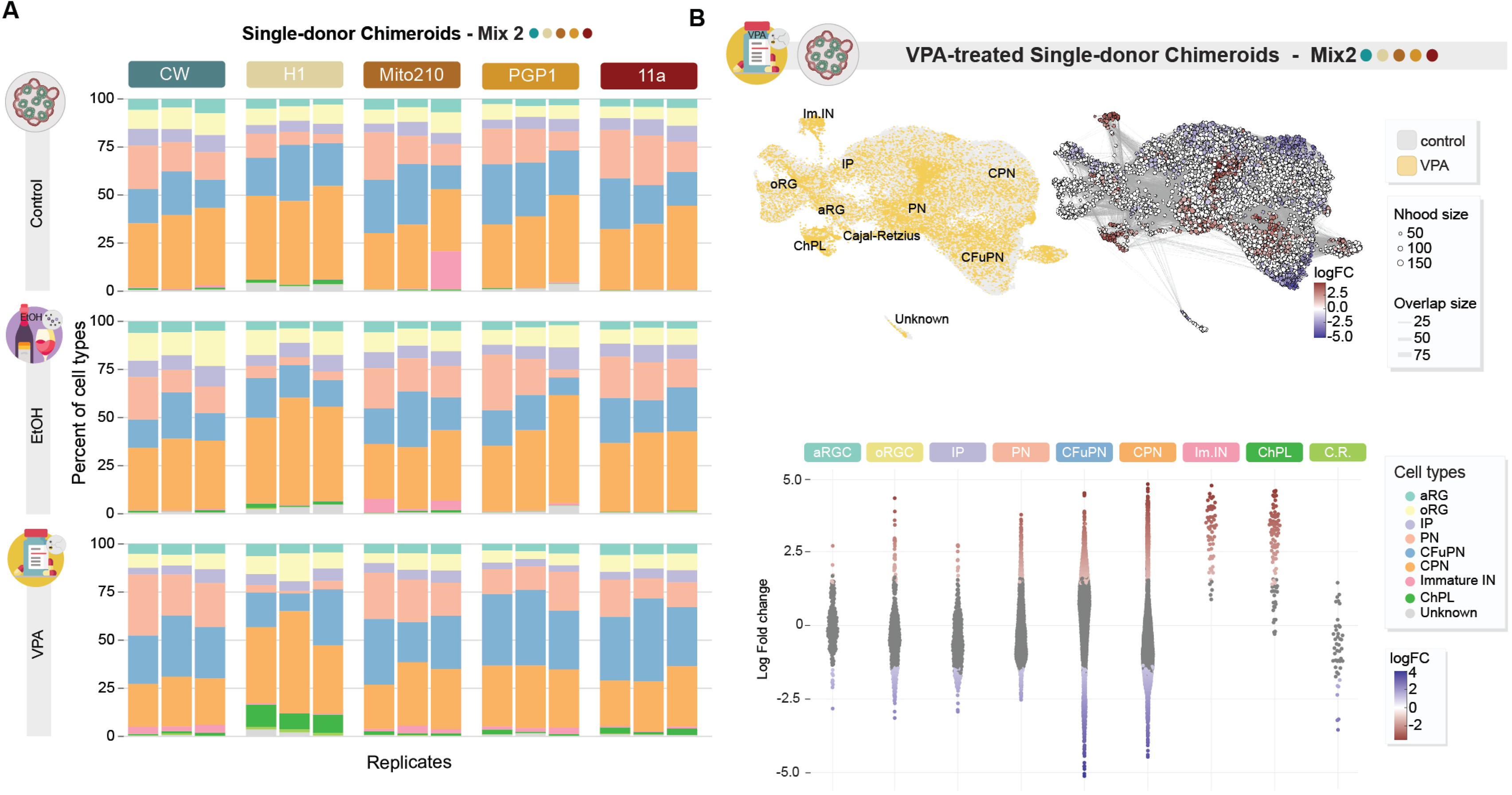
VPA and EtOH treatment cause alterations in the transcriptome profile of single-donor Chimeroids. **A,** Stacked barplot showing cell type composition for control and treated single-donor Chimeroids. **B,** Upper-left panel: cell-type specific changes in single-donor Chimeroids treated with VPA (yellow) vs control (grey). Upper-right panel: UMAP showing overlapping neighborhoods of cells, as calculated using Milo. Red and blue colors indicate neighborhoods with significant enrichment for VPA-treated cells or control cells, respectively. Point size indicates the number of cells in a neighborhood, and edge thickness indicates the number of cells shared between pairs of neighborhoods. Bottom panel: Beeswarm plot showing shifts in the composition of neighborhoods of cells in response to VPA treatment in single-donor Chimeroids, grouped by the cell-type identity of those neighborhoods. Each point represents a neighborhood of 50-200 cells with similar gene expression profiles. The vertical axis indicates the enrichment of VPA-treated cells within a neighborhood, with positive log-fold change values indicating more than expected VPA-treated cells, and negative values indicating fewer than expected VPA-treated cells. Neighborhoods are colored based on statistical significance of that enrichment: grey, not significantly different from random; red, significant over-enrichment; blue, significant under-enrichment. If most neighborhoods within a cell type collectively shift up or down, it implies an overall gain or loss, respectively, of that cell type in VPA-treated Chimeroids. Cell types with neighborhoods that form long tails of both over-enrichment and under-enrichment are likely to have treatment-induced changes in expression profile, without necessarily changing in abundance.

## Supplementary tables

Supplementary Table 1: Set of donors used in each mix used to create Chimeroids.

Supplementary Table 2: Marker Gene Expression Correlation related to Fig. 3c.

Supplementary Table 3: Post-hoc pairwise comparisons in cell-type proportions split by donors in Fig. 5c.

Supplementary Table 4: List of antibodies used in this study.

